# Decoding UTRs by applying explainable AI to a genomic foundation model

**DOI:** 10.64898/2025.12.04.692303

**Authors:** Logan Brase, Declan R. Creamer, Yuliya Shapovalova, Mark P. Ashe, Hilary L. Ashe, Magnus Rattray

## Abstract

**Background:** The regulation of mRNA decay and translation is crucial for cellular function and development; however, the complex interplay of RNA-binding proteins (RBPs) regulating these processes remains incompletely understood. Recent advances in genomic foundation models present new opportunities for decoding the regulatory grammar embedded within mRNA untranslated regions (UTRs). Here, we leverage explainable artificial intelligence to systematically identify RBP motifs that influence translation and mRNA decay during *Drosophila melanogaster* development.

**Results:** We extended the training of GENA-LM Fly, a genomic foundation model, on all 5’ and 3’ UTR pairs from the *D. melanogaster* genome. We separately fine-tuned it using ribosome density (RD) and mRNA decay (half-life) data from the embryonic maternal-to-zygotic transition (MZT). Using SHapley Additive exPlanations (SHAP) analysis, we identified the sequence regions most influential for prediction and performed motif enrichment analysis to discover associated RBP binding sites. We identified 42 unique RBPs associated with increasing (n=23; e.g., Rnp4f, Mxt) or decreasing (n=19; e.g., Aret/Bruno, Rox8, Sxl, Orb2) RD and 18 unique RBPs associated with increasing (n=6) and decreasing (n=12; e.g., Cnot4, Rbp9, Rox8) mRNA half-life. Using publicly available PAR-CLIP data, we validated our Orb2 signal in a *Drosophila* cell line. Furthermore, feature ablation and shuffling experiments revealed the contributions of different sequence components to model performance. Our approach significantly outperformed naïve high-versus-low RD comparisons, demonstrating the power of model explainability in biological discovery.

**Conclusions:** This study demonstrates that genomic foundation models, when combined with explainability methods, can discover meaningful biology even without drastically improving the underlying prediction accuracy. The identified RBP motifs provide new insights into post-transcriptional regulatory elements that govern RD and decay during early development.

## Background

During early embryonic development, precise control of the rate of protein synthesis enables rapid changes in protein abundance without corresponding changes in transcription rates (1,2). This regulatory layer is in part mediated by RNA-binding proteins (RBPs) that recognize specific sequence motifs within the 5’ and 3’ untranslated regions (UTRs) of target mRNAs (3). Controlling the decay rate of individual transcripts is another mechanism that regulates functional state and development (4) and is also influenced by the binding of RBPs to the 5’ and 3’ UTRs.

Despite decades of research, our understanding of the complex regulatory grammar governing translation remains incomplete. Traditional approaches to dissect protein synthesis have relied on techniques such as polysome profiling, ribosome footprinting, reporter assays, and biochemical labelling to generate data for the calculation of ribosome density (RD; RD is frequently used as a proxy for translation efficiency in the literature) or on the perturbation of individual RBPs(5–8). While these strategies have provided valuable mechanistic insights, they are typically constrained to one factor or sequence feature at a time and are not well suited to capturing the combinatorial and context-dependent nature of post-transcriptional regulation.

The challenge of capturing these combinatorial relationships in early statistical models, which relied on limited, precomputed features, led to the application of deep learning architectures that could process sequences as inputs, such as convolutional neural networks (CNNs) and recurrent neural networks (RNNs). Models such as the multitask CNN RiboNN (9) for RD and the hybrid network Saluki (10) for mRNA half-life achieved state-of-the-art performance on mammalian systems. However, these architectures can still fail to capture the full scope of long-range dependencies such as those between the 5’ and 3’ UTRs, and equivalent models for *Drosophila* are lacking.

The recent emergence of genomic foundation models represents a paradigm shift in computational biology driven by the need to model these long-range interactions. These models, pretrained on vast genomic sequences, capture the patterns and underlying syntax in DNA and RNA and can be fine-tuned for a variety of downstream tasks, where they have demonstrated a remarkable ability to capture complex sequence patterns and regulatory relationships (11–15). Notably, the recent transformer model FlashRNA (15), trained on the same datasets as Saluki and RiboNN, outperformed both architectures in predicting mRNA half-life and ribosome occupancy, further establishing transformers as the dominant approach for modelling long-range and context-dependent regulatory information.

Although the perceived "black box" nature of these models has limited their use for biological discovery, the application of explainability methods can make them a powerful tool for uncovering regulatory sequence features. As the name suggests, explainable AI aims to explain the inner workings of machine learning models and map influence scores to specific input features. Several of these methods have been applied to transformer models (16–19). Of these, SHAP is widely used because of its model-agnostic nature and theoretical guarantees (16), making it appealing even if it is not the most computationally efficient algorithm.

Conceptually, SHAP estimates how much each nucleotide contributes to the model’s prediction by comparing the prediction made with that nucleotide present to predictions made when it is replaced with a neutral baseline. These comparisons yield a quantitative score that reflects whether a given region increases or decreases the final model prediction. SHAP therefore provides a principled way to highlight the UTR sequence segments key to translation regulation, enabling downstream motif enrichment analyses to identify candidate RNAIZlbinding proteins associated with these processes.

In this study, we combine a genomic foundation model with explainable AI to systematically identify RBP motifs that are predicted to regulate RD and mRNA decay via the UTRs during *Drosophila* development. To clarify, our aim is not to produce a state-of-the-art model, but to apply explainable AI to an existing model to extract biological hypotheses relating to the UTRs. Our approach leverages SHAP analysis to highlight the sequence regions most influential for prediction, followed by motif enrichment analysis to identify associated RBP binding sites. This methodology provides a powerful framework for decoding the regulatory sites governing post-transcriptional control.

## Results

### Genomic foundation model successfully predicts ribosome density

We successfully extended the training of the GENA-LM Fly(12) genomic foundation model on *Drosophila* UTR sequences and fine-tuned it on RD data (Figure 1). The extended pretraining phase incorporated the 5’ and 3’ UTR sequence pairs from 30,284 transcripts (21,518 train; 4,494 val; 4,272 test), enabling the model to learn sequence representations specific to *Drosophila* regulatory elements (GENA-LM Fly UTR, Table S1). Fine-tuning on 6,019 transcripts (4,213 train; 901 val; 905 test) with reliable RD measurements combined with extra features (codon frequencies, GC content, sequence lengths, and mean free energy (MFE; proxy for secondary structure)) yielded strong predictive performance across train, validation, and test sets (FlyUTR-RD, Figure 2a, Figure S1, Table S2).

**Figure 1.**
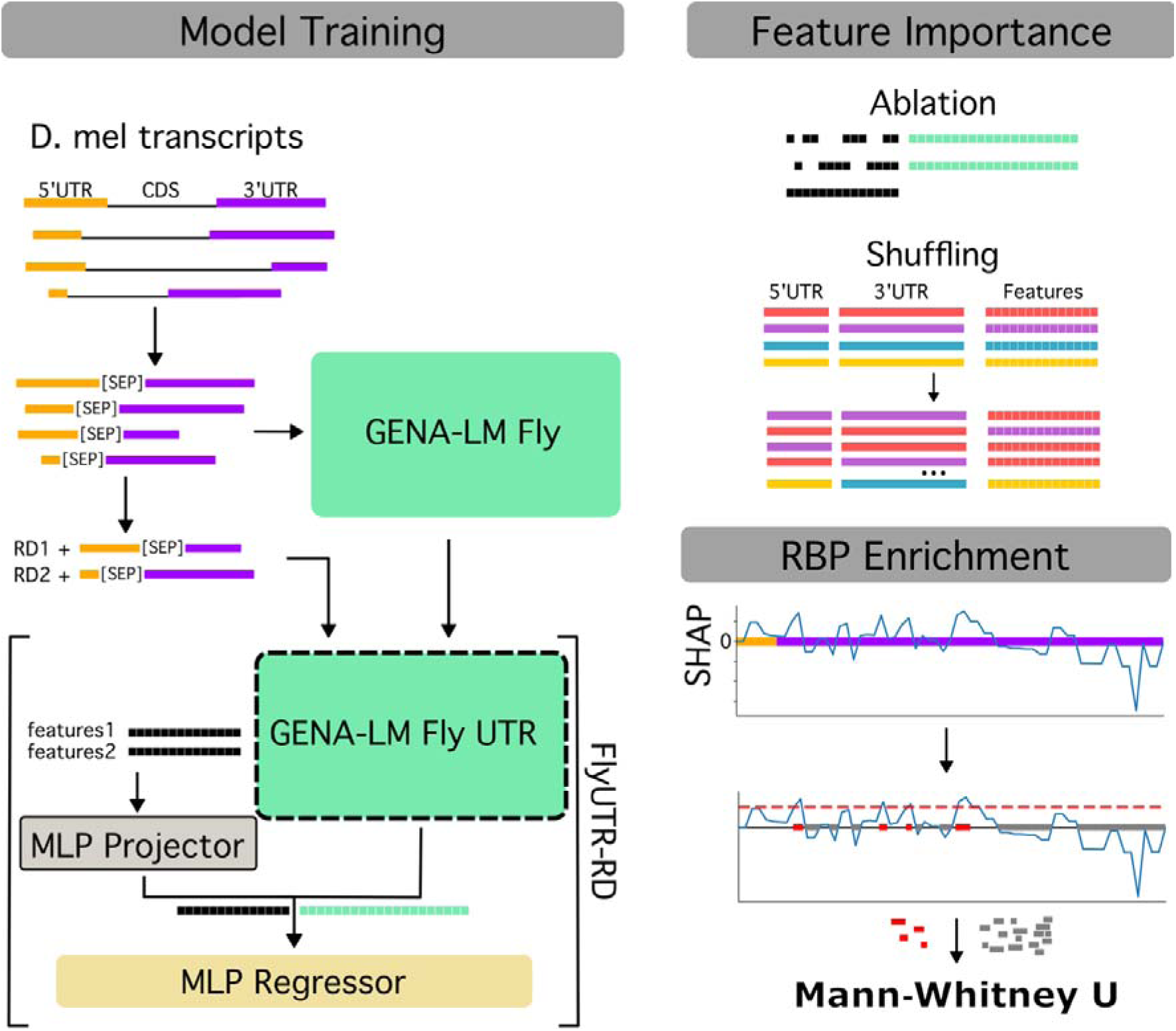
Study overview. Model training: Schematic of extended model training, fine-tuning, and final model structure. The GENA-LM Fly foundation model was subjected to extended pre-training using masked learning on D. melanogaster UTR sequence pairs to generate the GENA-LM Fly UTR model. To predict ribosome density (RD), a multilayer perceptron (MLP) projector and regressor were integrated with the foundation model, and the full architecture (FlyUTR-RD) was trained on transcripts with reliable RD measurements and additional features (e.g., codon frequencies and sequence lengths). Feature importance: Schematics of feature ablation and shuffling to identify the impact of features on the final model’s performance and prediction. RBP enrichment: Schematic for leveraging explainable AI to identify RBP motif enrichment in transcript regions with high absolute SHAP scores using Mann-Whitney U tests.

**Figure 2.**
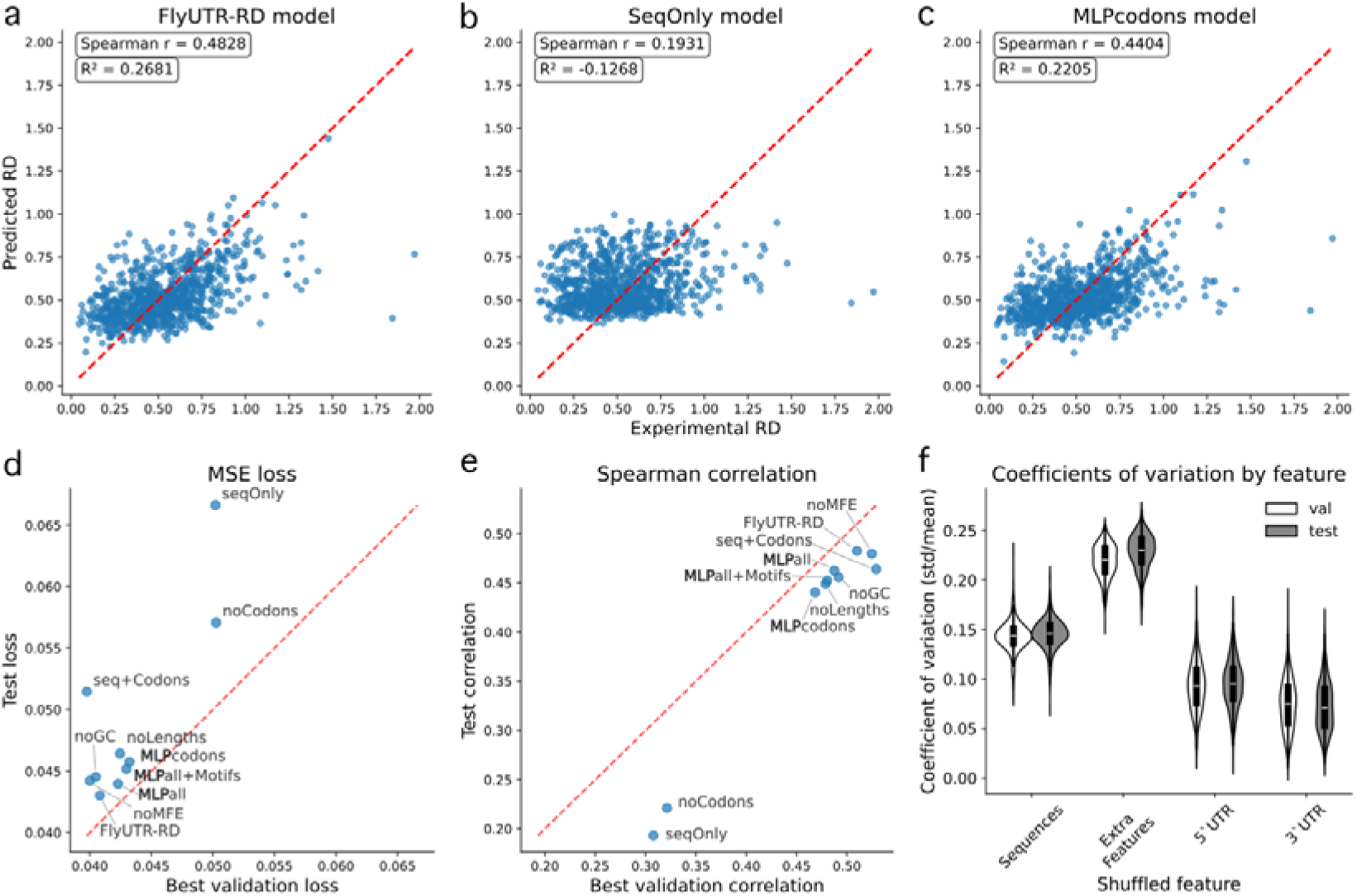
Model performance and feature importance analyses. a-c) Scatter plots of experimental RD versus predicted RD for the FlyUTR-RD model using the test data (a), the seqOnly model (b), and the MLPcodons model (c). d,e) Scatter plots of the validation and test losses (e, lower is better) and Spearman coefficients (e, higher is better) for FlyUTR-RD and ablation models. Bayesian optimization was used to optimize the hyperparameters for each model. Models starting with “MLP” are multilayered perceptron models as opposed to the fine-tuned genomic foundation models. Descriptions for each model can be found in the Feature ablation section of the Methods. f) The coefficient of variance for each transcript when shuffling each feature group in the FlyUTR-RD model using the validation (white) and test (gray) datasets. Sequences: shuffling of UTR pairs while keeping all else the same, ExtraFeatures: shuffling of matching lengths, GC content, MFE, and codon frequencies while keeping the UTRs the same, 5’UTR: shuffling of just the 5’ UTR, 3’UTR: shuffling of just the 3’UTR. A higher coefficient of variation indicates greater influence on prediction.

We included codon frequences instead of the full coding sequence (CDS) for two reasons. (1) The GENA-LM model has a context limit of 512 tokens which prohibits the inclusion of full-length transcripts. As we were most interested in the UTRs, we decided to remove the CDS. (2) Many post-transcriptional regulation processes are protein class specific and could be inferred from protein domains in the CDS. We want the model to learn features encoded in the UTRs not correlations with protein classes.

### Feature ablation validates model architecture choices

We created several fine-tuned models each with different combinations of sequence and features (feature ablation), to identify the best performing model and highlight the importance of different features for prediction. The model containing the sequence and all extra features (FlyUTR-RD) achieved a Spearman correlation of ρ = 0.48 between the predicted and experimental RD values on the test set (Figure 2a, d-e, Table S2). FlyUTR-RD’s performance is consistent with the spearman correlations of early mammalian models (UTR-LM, Optimus, Cao-RF, Kipoi, Mtrans, RNAFM, and RNABERT) compared in an RD benchmarking analysis in three different tissues (muscle: median ρ = 0.48, PC3: median ρ = 0.45, HEK: median ρ = 0.47) (20). Later mammalian models like Saluki have stronger performance, but this is mostly achieved by improved training datasets.

FlyUTR-RD outperformed both the model containing only the sequences with no extra features (seqOnly, ρ = 0.19, Figure 2b, d-e, Table S2) and a multilayer perceptron (MLP) model trained only on codon frequencies with no sequence or other extra features (MLPcodon, ρ = 0.45, Figure 2c, d-e, Table S2). These results support the current view that RD is largely determined by codon frequency (21,22), but also confirm that UTRs further fine-tune the RD value, as shown by the slight increase in model performance after UTR sequence inclusion. This fine-tuning is consistent with the known function of UTRs via RBP and microRNA binding.

Transcript length and GC content helped the model generalize, as we can see that the model without them (seq+Codons) had the best validation loss and correlation, but did not perform as well on the test set (Figure 2d,e). Furthermore, length seems to have a greater impact on performance than GC content. This could be driven by an unmodeled connection between UTR length and UTR features that regulate mRNAs such as untranslated open reading frames (uORFs), hairpins and loops. The model without MFE (noMFE) comes in second to FlyUTR-RD in both the loss and the correlation on the test set indicating that MFE has the least effect on model performance (Figure 2d,e). We postulate that the dynamics of RNA secondary structures and their interactions have a greater impact than can be captured in a single MFE score. FlyUTR-RD was used in all of the downstream RD analyses.

Feature shuffling experiments using FlyUTR-RD revealed that the extra features, driven by the codon frequencies, have the largest coefficients of variation (greater impact on model predictions) as expected. They also show that the 5’ UTR sequences have higher coefficients of variation than the 3’ UTR sequences, suggesting that the 5’ UTR is generally more important in defining transcript RD (Figure 2f). Interestingly, the shuffling of paired UTR sequences has a tighter distribution than when the 5’ and 3’ are independently shuffled. This could suggest that the transformer architecture is capturing interactions between the 5’ and 3’ UTRs; that the UTR pairs have been evolutionarily optimized together potentially in connection to the “closed-loop” model of translation initiation (23).

Although the ablation analysis demonstrates the improved performance of FlyUTR-RD over the MLP models, its real strength becomes apparent when the model is interrogated to discover the learned patterns from the UTR sequences.

### SHAP analysis reveals position-specific regulatory patterns

SHAP explainability analysis using FlyUTR-RD identifies distinct positional patterns of regulatory importance within the 5’ and 3’ UTRs (Figure 3). In the 5’ UTRs, regions immediately downstream of the 5’ cap presented consistently large positive SHAP values, indicating their importance in promoting RD (Figure 3a). This aligns with our shuffling analysis and the current understanding that ribosomes are recruited to mRNAs through translation factors such as eIF4E and eIF4G, which bind to the mRNA cap structure (24). Additionally, the proximal end of the 5’ UTR shows a further decrease in positive SHAP values, whereas the negative values remain consistent across the UTR. The 3’ UTR shows consistent values for both the positive and negative SHAP scores, except for a decrease in the magnitude of positive scores at the distal end (Figure 3b). The positive scores are generally higher in the 5’ UTR than in the 3’ UTR, suggesting a greater role for the 5’ UTR in regulating RD. The negative magnitudes are consistent across both UTRs; this could indicate that factors decreasing RD (e.g., RBP binding) are uniformly distributed across both UTRs rather than localizing to a specific region of one of the UTRs.

**Figure 3.**
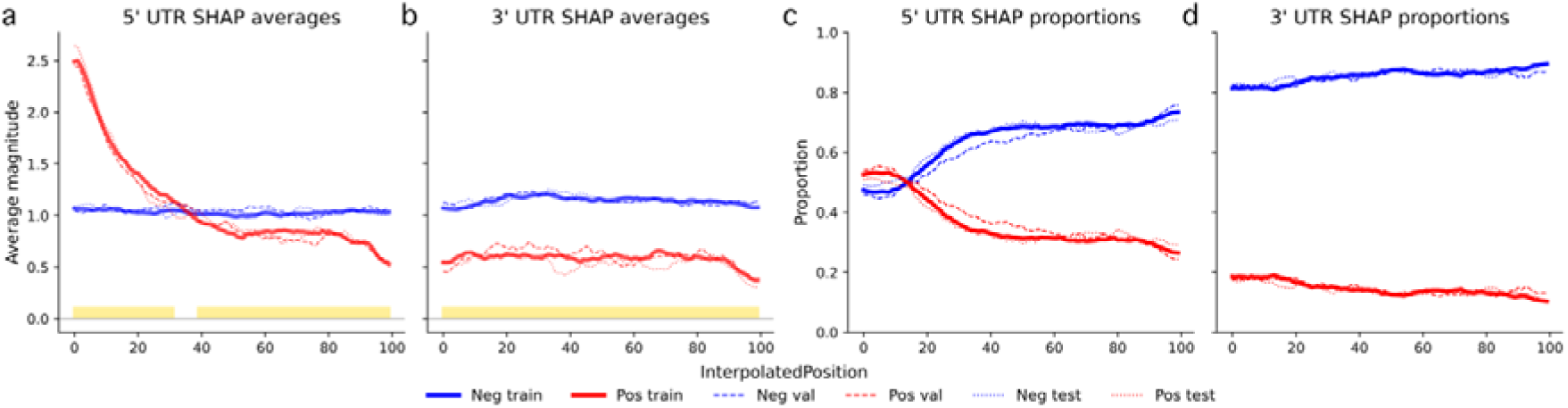
Global SHAP patterns for RD prediction across UTRs. a, b) The mean positive (red) and negative (blue) SHAP scores for the train, validation, and test sets across the normalized lengths of the 5’ UTRs (a) and 3’ UTRs (b). A linear model was fit at each position along the corrected lengths of the UTR regions. The model is defined as (SHAP ∼ sign + dataset), where sign is 1 or 0 for positive and negative SHAP, respectively, and dataset is train, validation, or test. Yellow bars at the bottom of the plot indicate a significant difference (BenjaminiUHochberg corrected) between the positive and negative SHAP magnitudes at that position as determined by the linear model. c, d) The proportion of transcripts that have a positive (red) or negative (blue) SHAP value at that position in the 5’ UTR (c) and 3’ UTR (d). The train, validation, and test sets are plotted separately.

In both UTRs, there are proportionally more negative SHAP values than positive (Figure 3c, d). The exception to this is at the very beginning of the 5’ UTRs, where there are more positive scores. This suggests that the specific control of ribosome recruitment at the 5’ UTR is a core component of overall mRNA translation. Although both UTRs generally have more negative values, the 3’ UTR consistently has greater than 80% negative values at each position, whereas the 5’ UTR peaks at less than 80% negative values. This also points to a greater impact from the 5’ UTR on increasing RD. Furthermore, the proportion of negative scores gradually increases from the proximal end of the 3’ UTR to the distal end. This could indicate that the model is capturing the impact of 3’ UTR length on RD directly from the sequences in addition to its inclusion as an extra feature. These positional patterns were consistent across train, validation, and test datasets (Figure 3c, d), indicating robust model behavior.

### Motif enrichment analysis identifies key regulatory RBPs

We individually tested the 5’ and 3’ UTR regions for motif enrichment by comparing the MAST (25) calculated RBP counts in the high-magnitude SHAP regions to those in the control regions using Mann□Whitney U tests. We considered RBPs as hits if they were significant in the train set and if the enrichment direction was consistent across the train, validation, and test sets (Figure 4, Figure S2). There are 20 and 17 RBPs associated with increased RD in the 5’ and 3’ UTRs, respectively, with 14 overlapping (Figure 4a,b; Table 1, Figure S3). A total of 19 and 17 RBPs are associated with decreasing RD in the 5’ and 3’ UTRs, respectively, with 17 overlapping (Figure 4c,d; Table 1, Figure S3). Notably, Sf2, Mxt and Rnp4f are linked to translational activation (26–29), and A2bp1 (Rbfox1), Aret (Bruno), Bru-3, Orb2, Pum, Rbp9, Rox8, and Sxl are known translational repressors (30–38), validating our analytical approach (Table 1).

**Figure 4.**
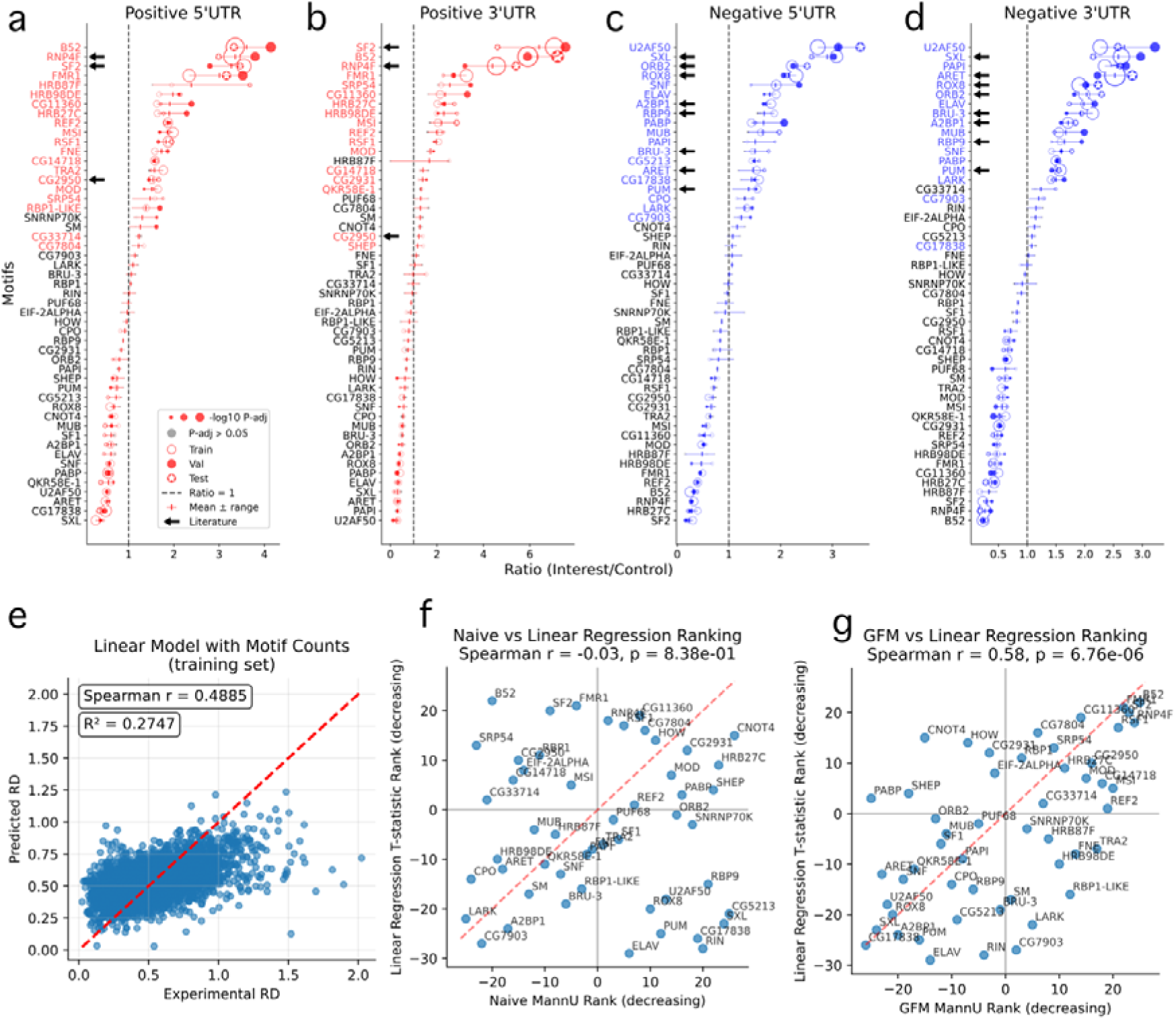
Motif enrichment results. a-d) Dot plots of the ratio of mean RBP counts in the high-magnitude SHAP regions vs the control regions for motifs from the train, validation, and test sets split by high (a, b) and low (c, d) SHAP scores and 5’ (a, c) and 3’ (b, d) UTRs. The error bars represent the minimum and maximum values for each motif across the train, validation, and test sets. The names of RBPs that are significantly enriched in the train set and have a consistent direction of effect in the validation and test sets are colored. Significant RBPs that have support from the literature are marked with a black arrow (see Discussion). e) Scatter plot of experimental RD vs RD predicted by the linear model trained on all the extra features plus motif counts per transcript (trained and predicted on the training set only). f, g) Scatterplots comparing the rank of the linear regression T-statistic to the rank of the signed p-values from the MannUWhitney U motif enrichment results for the high RD sequences from the ‘naïve’ high vs low model (f) and the positive high-SHAP regions from FlyUTR-RD (g). Positive values indicate the RBP increases the RD predicted value in the linear model and is enriched in the Mann-Whitney U analysis. Negative values indicate the RBP decreases the RD predicted value in the linear model and is depleted in the Mann-Whitney U analysis.

**Table 1.**
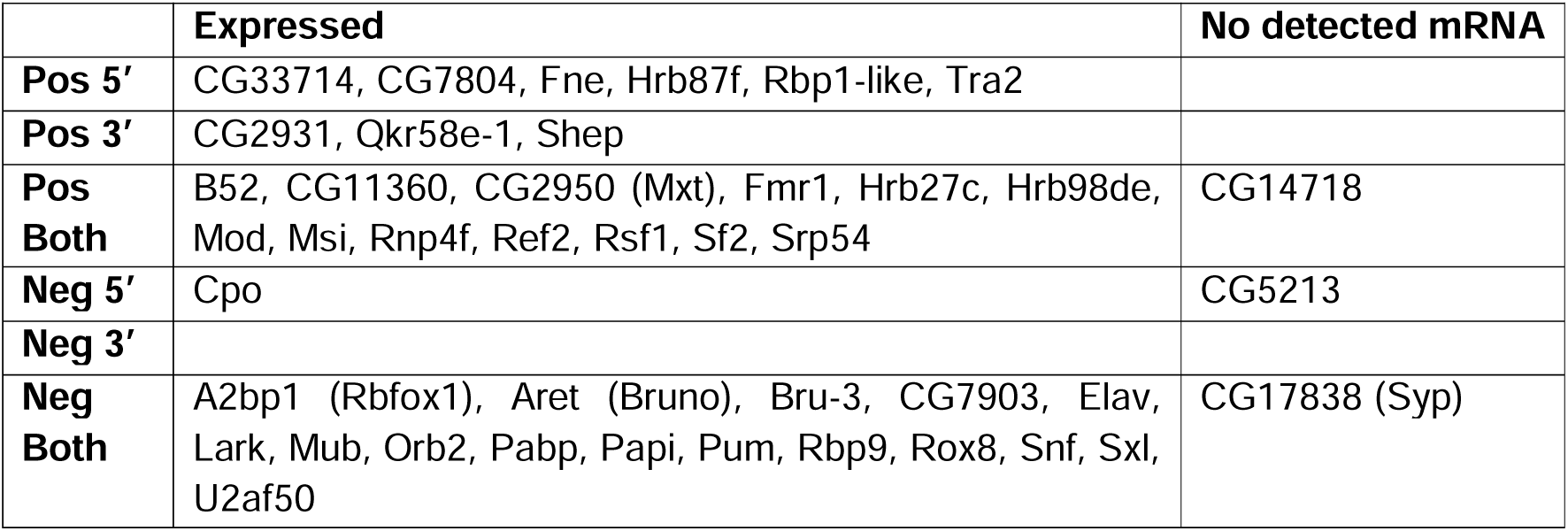
RBPs enriched in high-magnitude SHAP regions by UTR. Pos = regions associated with increasing ribosome density, Neg = regions associated with decreasing ribosome density.

Using the Ribo-seq and RNA-seq results used for RD calculation, we confirmed that all but three of these RBPs had detectable levels of mRNA at this stage of development (Table 1). (We note, however, that mRNA levels do not directly correlate with protein abundance, so the lack of detectable mRNA does not confirm the absence of the RBP.)

The motif enrichment results showed remarkable consistency across train, validation, and test sets, with the validation and test sets agreeing in the effect direction for all but five RBP motifs (Pos 3’: Hrb87f, Tra2; Neg 3’: Rin, Cpo, CG5213) significantly enriched in the training sets (Figure 4a-d, Figure S3). Furthermore, these five nonconforming RBPs either have low significance and/or low effect size in the training sets. This consistency demonstrates the robustness of our findings and suggests that the identified RBPs represent genuine regulatory factors rather than dataset-specific artifacts.

### Model-based approach outperforms naïve analysis

To validate the superiority of our explainability-based approach, we compared our results with a naïve differential motif enrichment analysis that directly compared the UTRs of the high- and low-RD transcripts. The naïve approach was severely underpowered with no significant hits. We also ran a linear regression model to predict RD using all the extra features and motif counts within each transcript. We anticipated that the linear model would lack the power to detect significant associations with motifs, but would generally capture effect directions correctly, thereby providing a baseline for comparison. As expected, the linear model was underpowered but identified one motif (Elav) as significantly associated with lowering RD (Figure 4e).

We then calculated the correlation between the motif enrichment ranks of the naïve analysis and the SHAP-based analysis using the positive 5’ UTR results from the FlyUTR-RD model to the ranks of the motif T-scores from the linear model. Not only did the naïve analysis fail to identify any significant motifs, but it also had no correlation with the linear model (ρ =-0.03, p-val = 8.38e-1, Figure 4f) indicating that even a simple linear model outperforms this approach. Conversely, the correlation between FlyUTR-RD and the linear model was quite strong (ρ = 0.58, p-val = 6.76e-6, Figure 4g). This confirms the accuracy of the SHAP explainability technique and highlights the added power FlyUTR-RD provides in identifying regulatory relationships. This superior performance likely reflects its ability to account for combinatorial effects and sequence context.

### PAR-CLIP validation of Orb2

To further validate that the model’s SHAP attributions reflect genuine binding activity, we sought to cross-reference the results with PAR-CLIP/CLIP-seq datasets. Of all the RBP hits, only Orb2 had publicly available PAR-CLIP data, however, it is generated from *Drosophila* S2 cells rather than the embryo at the MZT. Despite this limitation, we used this data to perform both a UTR- (hits anywhere in the UTR) and position-level (motif overlaps CLIP peak) enrichment analysis.

For the 5’ and 3’ UTRs independently, we defined three gene sets: Group A (no Orb2 MAST motif hit), Group B (Orb2 MAST hit present but not overlapping a low-SHAP window), and Group C (Orb2 MAST hit overlapping a low-SHAP window, i.e. model-predicted Orb2 targets). Group C genes were significantly enriched for CLIP targets compared to Group B for both 5’ UTR targets (67.9% vs 58.1%, OR=1.52, p=0.013, Figure 5a) and 3’ UTR targets (87.9% vs 82.3%, OR=1.56, p=0.019, Figure 5b), confirming that SHAP-guided selection of motif hits improves prediction of experimentally validated Orb2 targets beyond motif presence alone. The any-UTR comparison did not reach significance (88.9% vs 86.5%, OR=1.24, p=0.135, Figure 5c), consistent with the UTR-specific signal being diluted when considering both UTR types jointly.

**Figure 5.**
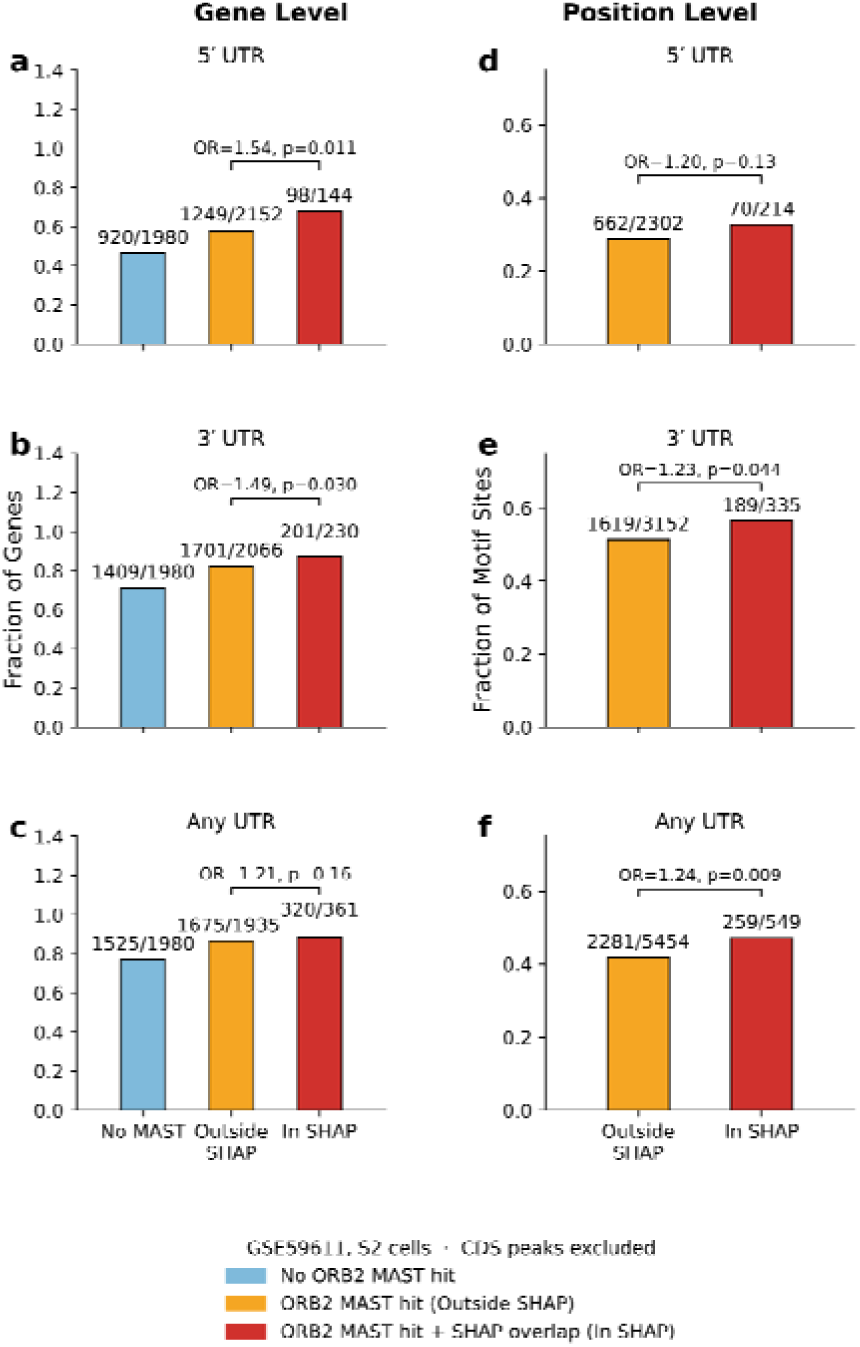
PAR-CLIP validates Orb2 associated genes and positions prioritized by SHAP. Left column (a-c), gene-level analysis: genes were classified as No MAST, Outside SHAP, or In SHAP, corresponding to no Orb2 MAST hit in the relevant UTR region, at least one Orb2 MAST hit without overlap with a low-SHAP region, or at least one Orb2 MAST hit overlapping a low-SHAP region, respectively. Right column, position-level analysis: unique genomic Orb2 motif sites were classified as Outside SHAP or In SHAP according to whether the motif fell outside or within a low-SHAP region. Rows show analyses performed 5′ UTR only (a,d), 3′ UTR only (b,e), and across merged UTR (c,f), hits. Bar heights indicate the fraction of genes or motif sites supported by PAR-CLIP, and numbers above bars indicate the number of PAR-CLIP-supported observations over the total number in that class. Brackets denote the Fisher’s exact test comparing SHAP-overlapping versus non-overlapping categories, with odds ratio (OR) and one-sided P value shown above each comparison. Colors denote no Orb2 MAST hit (blue), Orb2 MAST hit outside SHAP regions (orange), and Orb2 MAST hit overlapping SHAP regions (red).

At the position level, Orb2 motif sites falling within low-SHAP windows showed significantly higher CLIP peak overlap than sites outside these windows across both UTRs combined (47.4% vs 41.8%, OR=1.26, p=0.007, Figure 5f) and for 3’ UTR sites specifically (56.8% vs 51.3%, OR=1.25, p=0.034, Figure 5e), with a consistent but non-significant trend in the 5’ UTRs (32.9% vs 28.8%, OR=1.21, p=0.12, Figure 5d). Together, these results indicate that SHAP attributions from the FlyUTR-RD model prioritize biologically meaningful Orb2 binding sites at both UTR and binding site resolution. This is even more notable when considering that the data came from different sources (whole embryo vs S2 cell line).

### Fine-tuning for mRNA decay

After observing the success of this method for RD, we investigated whether it would also be useful on a much smaller mRNA decay dataset. Decay estimates were obtained for a subset of transcripts expressed zygotically, but not maternally, close to the MZT (39). We converted the decay rates to half-lives and log transformed the values for training. Starting with the same extended pretraining model, we fine-tuned for half-life using the same extra features but for the smaller dataset of 615 transcripts (430 train, 94 val, 91 test). After running the same ablation analyses performed on the RD model, we identified the model trained on sequences and all extra features except codons (noCodons model) as achieving the best correlation (ρ = 0.365, Figure 6a-c) with the experimental half-lives and used this model to visualize SHAP patterns and test for motif enrichment.

**Figure 6.**
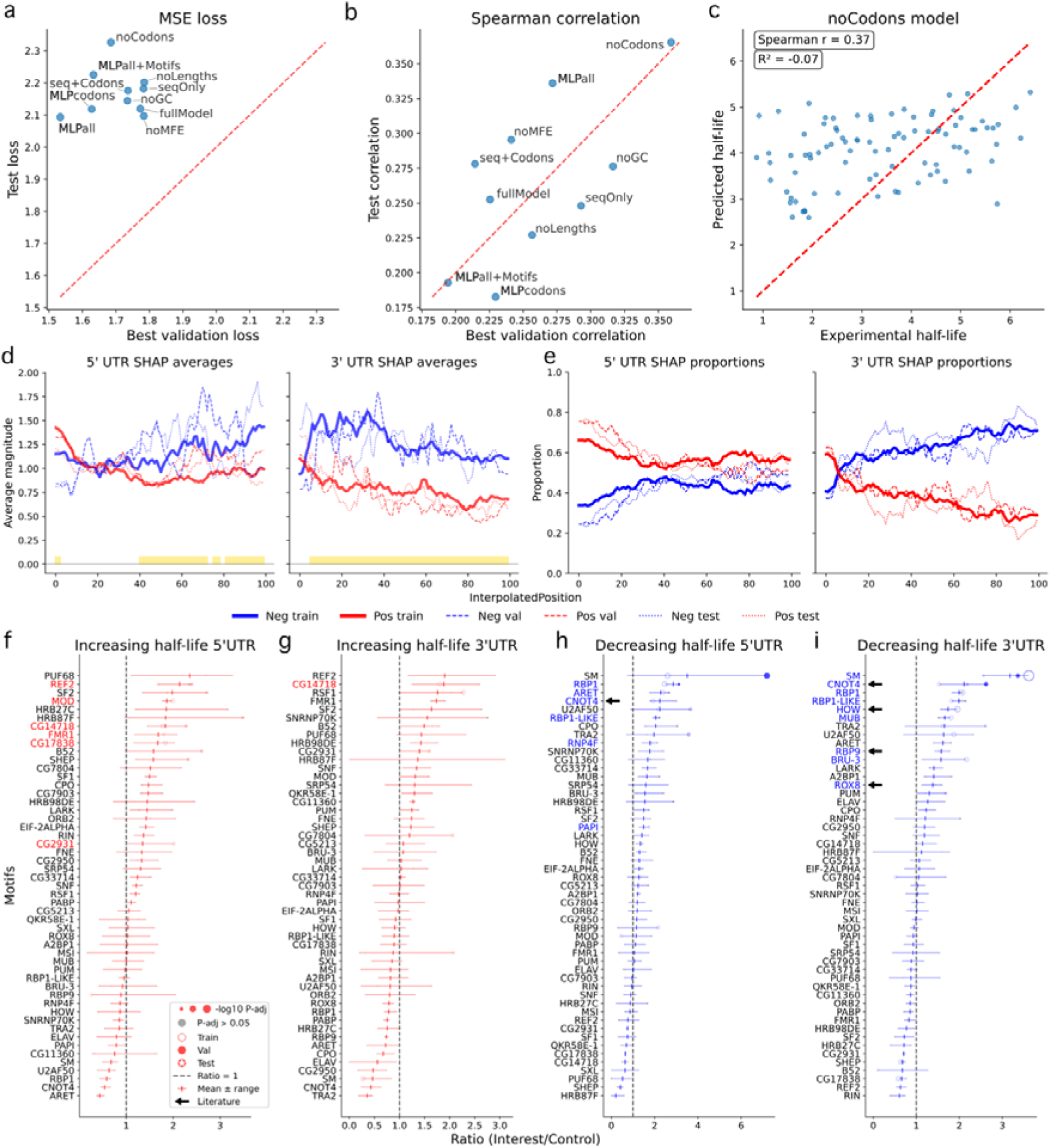
Half-life model performance, SHAP patterns and motif enrichment. a, b) Scatter plots of the validation and test losses (a, lower is better) and Spearman coefficients (b, higher is better) for FlyUTR-RD and ablation models. c) Scatter plot of the experimental versus predicted half-lives from the noCodons model for the test set. d) The mean positive (red; increasing half-life) and negative (blue; decreasing half-life) SHAP scores for the train, validation, and test sets across the normalized lengths of the 5’ UTR (left) and 3’ UTR (right). A linear model was fit at each position along the corrected lengths of the UTR regions. The model was defined as (SHAP ∼ sign + dataset), where sign is 1 or 0 for positive (increasing half-life) and negative (decreasing half-life) SHAP, respectively, and dataset is train, validation, or test. Yellow bars at the bottom of the plot indicate a significant difference (BenjaminiUHochberg corrected) between the positive and negative SHAP magnitudes at that position as determined by the linear model. f) The proportion of transcripts that have a positive (red) or negative (blue) SHAP value at that position in the 5’ UTR (left) and 3’ UTR (right). The train, validation, and test sets are plotted separately. f-i) Dot plots of the ratio of mean RBP counts in the high-magnitude SHAP regions vs the control regions for motifs from the train, validation, and test sets split by high (f,g) and low (h,i) SHAP scores and 5’ (f,h) and 3’ (g,i) UTRs. The error bars represent the minimum and maximum values for each motif across the train, validation, and test sets. The names of RBPs that are significantly enriched in the train set and have a consistent direction of effect in the validation and test sets are colored. Significant RBPs that have support from the literature are marked with a black arrow (see Discussion).

The global SHAP patterns highlight the importance of the distal end of the 5’ UTR and the proximal end of the 3’ UTR, as both have the highest magnitudes and proportions of positive SHAP values compared with the rest of their respective UTRs (Figure 6d,e). The features associated with decreasing half-life seem to be strengthened at the proximal halves of both UTRs apart from the very proximal end of the 3’ UTR, where the magnitudes and proportions are low for the negative SHAP values.

Despite the relatively weak predictive performance of the model, it still learned valuable patterns from the UTRs. The regions associated with decreasing half-life are enriched for Cnot4, Rbp1, and Rbp1-like binding sites in both UTRs with another three (Aret, Papi, Rnp4f) and six (Bru-3, How, Mub, Rbp9, Rox8, Sm) RBPs enriched in the 5’ and 3’ UTRs, respectively (Figure 6h,i). How, Rbp9, and Rox8 all have known roles in destabilizing mRNA, which validate our approach even in this weaker model (40–42). The regions associated with increasing half-life are enriched for CG14718 motifs in both the 5’ and 3’ UTRs (Figure 6f, g); however, no detectable levels of mRNA for this RBP were found at the MZT according to our RD dataset (Table 1). The 5’ UTR is additionally enriched for CG2931, Fmr1, Ref2, and Mod motifs. (One additional motif, CG17838 (Syp), is enriched in the 5’ UTR but lacks detectable mRNA.)

## Discussion

Our study demonstrates the power of combining genomic foundation models with explainability methods to decode complex regulatory mechanisms governing mRNA translation and decay. The identification of 42 RBPs associated with translation regulation and 18 RBPs associated with mRNA decay/half-life provides new insights into post-transcriptional control mechanisms during *Drosophila* development and validates several previously characterized regulators.

### Negative regulators of ribosome density

The 17 RBPs overlapping in both UTRs for association with decreased RD fall into different known functional groups (Figure 4c,d). A2bp1 (Rbfox1), Aret (Bruno), Bru-3, Orb2, Pum, Rbp9, Rox8, and Sxl play known translational repression roles (30–38). There is evidence that CG17838 (Syp), Lark, and Mub have both repressive and activating roles (43–45), suggesting that during the MZT, they primarily work as repressors. Pabp is a known translational activator via its role in the “closed-loop” complex (23); however, recent studies have also demonstrated that Pabp can mediate translational repression, indicating that the role of Pabp is more complex than originally thought (46,47). This could suggest that during the MZT, Pabp might perform a similar repressive role depending on the mRNA context. SNF and U2af50 are known splicing factors (48,49), Papi is involved in PIWI (piRNA pathway) for transposon silencing (50), and Elav triggers alternative polyadenylation (APA) (51) to change the 3’ UTR length of transcripts. Further investigation is needed to determine whether these known functions indirectly affect mRNA translation or if these RBPs have direct translational repression roles in addition to their known functions.

Using an Orb2 PAR-CLIP dataset from *Drosophila* S2 cells, we showed that the FlyUTR-RD model prioritized Orb2 motif sites that map to true Orb2 binding sites (PAR-CLIP peaks). Not only does this validate our Orb2 results and give credibility to the other hits, but it shows that these results at least partially generalize between the distinct states of the early embryo (2-2.5 hours post-fertilization) and S2 cells (20-24 hours post-fertilization).

### Positive regulators of ribosome density

The 14 overlapping RBPs associated with increasing RD in both UTRs also fall into different groups according to known functional roles (Figure 4a, b). Sf2, Rnpf4 and Mxt (CG2950) are associated with translational activation (26–29), thus validating the findings from our model. Sf2 is a shuttling SR protein, classically thought of as a pre-mRNA splicing factor (28). However, Sf2 has been found associated with the translational machinery in the cytoplasm and to enhance translation via suppression of 4E-BP activity (52). Like Sf2, B52 and Srp54 are also shuttling SR proteins (52). Although we did not find previous analyses linking B52 and Srp54 to translation, many of the SR family of proteins are known to shuttle between the nucleus and cytoplasm where they regulate translation (27,53). Therefore, a similar function is probable for B52 and Srp54. Little is known about Ref2 other than its predicted role in exporting mRNA from the nucleus (53). Our results suggest that Ref2 may contribute to translational activation. Alternatively, the transcripts exported by Ref2 may generally exhibit higher RD values, leading to the observed association in our model. Fmr1, Hrb27c, and Msi are associated with both activating and repressing translation, which, according to our model, could mean that during the MZT, they predominantly function as activators. To our knowledge, CG14718 and Hrb98de only have connections to translational repression rather than activation as predicted in our study (54,55). Rsf1 is known for its role in mRNA splicing (56). This suggests that its splice variants may increase RD during development or it functions via a yet unknown mechanism. Additional studies are needed to unravel this further.

All hits that are only associated with one UTR have much weaker ratios and p-values, but two RBPs of note are Tra2 and Fne. Tra2, like other hits is known for its splicing function (57), but it is unclear if splicing is indirectly influencing RD or if it has direct connections to translational activation as well. Fne is in the same family as Elav and Rbp9 and is predominantly known for its role in neuronal development, even rescuing some functions in the loss of Elav (58). Being in the same family, Fne, Elav, and Rbp9 have very similar binding sites, yet only Fne is associated with increasing RD. As previously mentioned, Rbp9 has been linked to direct translational repression function, suggesting that Fne and Elav may have similar, yet undiscovered, roles in translational activation/repression.

### Regulators of mRNA half-life (decay)

The half-life dataset was much smaller than the RD dataset and therefore had less power, with a total of 18 unique RBPs associated with half-life (Figure 6f-i). Rbp9, Rox8, and How are associated with decreasing half-life via the 3’ UTR and have previously been shown to destabilize or accelerate the decay of specific transcripts (40–42). Cnot4 was linked to decreasing half-life in both UTRs. It is a nonessential part of the CCR4-NOT complex that catalyses deadenylation, a critical part of destabilizing mRNA for decay (59). This suggests that during the MZT, Cnot4 could function via this complex to accelerate degradation. The other 14 RBPs have no direct links to decay in the literature, but several act as translational repressors/activators (31,32,43,60,61), splicing factors (27,45,62), or components of the PIWI pathway (50,63).

Remarkably, our genome foundation model directed motif enrichment analyses have identified many of the well-characterized regulators of mRNA translation and stability in a non-biased manner, validating our analytical approach. Furthermore, it has identified many novel candidates revealing previously unknown components of regulation.

### SHAP explainability

The use of SHAP explainability provides quantitative importance scores that enable rigorous statistical analysis of motif enrichment, going beyond simple transcript level presence/absence comparisons. The superior performance of our model-based approach compared with that of naïve high-versus-low comparisons (Figure 4f, Figure S4) highlights the importance of accounting for sequence context and combinatorial effects in regulatory analysis. The distinct positional patterns of SHAP importance within 5’ UTRs (Figure 3) provide new insights into the architectural principles governing translation regulation. The increased strength and concentration of positive regulatory signals near the 5’ cap are consistent with the known importance of cap-dependent translation initiation (24).

We note that the token used as the baseline for the SHAP analysis is important. By default, the SHAP package for explaining transformers uses the [MASK] token as the baseline (16). We saw that this can result in imbalanced SHAP values (i.e., almost all positive SHAP scores). Using the token with an embedding closest to the average of all the embeddings mitigated this issue (see Methods).

We chose to map known RBP motifs onto our sequence regions highlighted by SHAP to make connections to known biology straightforward. As performed in other studies (9,64), these sequence regions could have been used to directly compute motifs, which would have facilitated the discovery of novel motifs rather than relying on known biology. Other tools modelling RD from sequence context, focus on evolutionary constraint (65), the Kozak sequence (20), or other features rather than on RBPs (9). Our study uniquely prioritizes known RBPs for functional validation as translational activators and repressors.

### Limitations and future directions

While our study provides valuable insights into the regulation of mRNA translation and degradation, several limitations should be acknowledged. First, our analysis focuses exclusively on the maternal-to-zygotic transition in *Drosophila*, and the identified regulatory relationships may not generalize to other developmental stages or organisms. However, we do note the consistency between our embryo data and the S2 cell line. Second, the correlation between SHAP importance and functional significance, while biologically plausible, has not been experimentally validated. This validation through targeted mutagenesis and reporter assays will be crucial for confirming the functional relevance of predicted regulatory elements. Future studies should extend this approach to additional developmental stages and species to assess the conservation and context dependence of the identified regulatory relationships.

Our analysis of degradation involved a small number of zygotically activated transcripts with consequently less power to detect informative sequence features. One factor driving the predictive success of models in the mammalian space is the high volume of available datasets. RiboNN leveraged 3,819 ribosomal profiling datasets, and Saluki calculated consensus half-life measurements from a meta-analysis of 39 human and 27 mouse datasets. Future studies producing and aggregating these datasets in *Drosophila* are needed to produce models as accurate as those in mammals.

Genomic foundation models facilitate critical knowledge transfer to data-scarce domains like RD and decay, yet the added extra features lack this pre-trained advantage, remaining bottlenecked by fine-tuning sample sizes. Our inclusion of codon frequencies, GC content, and MFE represents a pragmatic trade-off between context length, architectural complexity, and training data scale. Omitting the full or proximal CDS (first 30–60 nucleotides) likely misses key UTR-CDS interactions and the full scope of translation initiation. Furthermore, while global MFE proxies stability, it lacks the resolution of localized structural features; however, integrating spatially aware MFE values with transcript sequences exceeds the scope of standard foundation model fine-tuning. There is also a host of additional biological signals including Kozak sequences, dinucleotide frequencies, amino acid frequencies and secondary structure features like loop or hairpin counts that could improve model performance if properly balanced with sufficient training set size.

### Conclusions

We have demonstrated that genomic foundation models, combined with explainability analysis, can effectively decode regulatory mechanisms governing mRNA translation and decay during development. Our approach identified 42 RBPs with significant associations with translation regulation and 18 with associations to mRNA half-life, including both known regulators and novel candidates for future investigation.

This study establishes a general framework for applying explainable AI to regulatory genomics problems. As foundation models continue to grow in size and sophistication, explainability methods will become increasingly important for extracting biological insights from these complex systems, accelerating our understanding of gene regulation and its role in development, disease, and evolution.

## Methods

### Model architecture and training

We used GENA-LM Fly (12), a transformer-based genomic foundation model pretrained on genomic sequences from 298 drosophilid species, as the base architecture for our model. All training and evaluation were performed on transcripts grouped into ortholog clusters as defined by the DrosOMA database (https://drosoma.unil.ch/oma/home/). These groups, rather than individual transcripts, were then randomly split into training, validation, and test sets to minimize data contamination from highly similar sequences.

#### Extended pretraining

To adapt the base model to untranslated regions (UTRs), we performed extended pretraining using a masked language modelling objective (pytorch v2.5.0, pytorch-cuda v12.4, transformers v4.48.3). Specifically, we extracted all 5’ and 3’ UTR sequence pairs from the *Drosophila melanogaster* reference genome (BDGP6, GCA_000001215.4) and the corresponding Ensembl Rapid-Release gene annotation (GTFF: GCA_000001215.4, 2022_07). Sequences were tokenized using the byte-pair encoding (BPE) tokenizer provided with GENA-LM Fly, with the [SEP] token inserted between each 5’ UTR and its corresponding 3’ UTR. During training, 15% of tokens were randomly masked, and the model was trained to predict the masked tokens. Training was performed on Nvidia L40S GPUs.

#### Fine-tuning for ribosome density

Following extended pretraining, we fine-tuned the model on ribosome profiling data collected during the maternal-to-zygotic transition in *D. melanogaster*. For each transcript, we incorporated additional features: codon frequencies; GC content percentage; and length, which were calculated separately for the 5’ UTR, coding sequence (CDS), and 3’ UTR. We also included the mean free energy (MFE) for the full transcript (5’ UTR, CDS, 3’ UTR together) computed using linearFold (v1.0.1) (66) as a proxy for the secondary structure.

These features (n=68, FlyUTR-RD) were processed through an MLP projector to align their scale with the transformer’s output embeddings. The projector architecture consists of a linear layer (input dimension equal to the number of extra features, output dimension equal to the number of extra features), followed by ReLU activation, dropout, and a second linear layer of the same size. The projector output (n=68, FlyUTR-RD) was concatenated to the transformer’s hidden state representations (n=768) and fed into an MLP regressor for RD prediction. The regressor architecture included a linear layer (input dimension equal to the GENA-LM hidden size plus the number of extra features (n=836, FlyUTR-RD), output dimension of 256), followed by ReLU activation, dropout, and a final linear layer (input dimension of 256, output dimension of 1). The training objective was to minimize the MSE loss.

#### Hyperparameter optimization and training procedure

Model hyperparameters were optimized using Bayesian optimization via the Ray Tune (67) (v2.48.0) library with HyperOpt (68) (v0.2.7), running 100 trials with an optimization objective of maximizing the Spearman correlation on the validation set as this helped avoid fitting the mean. The training objective remained minimizing the MSE loss. Additional Ray Tune parameters include:

- ray_tune_max_epochs: 50
- ray_tune_initial_points: 5
- ray_tune_grace_period: 4
- ray_tune_reduction_factor: 2
- ray_tune_cpu_per_trial: 2
- ray_tune_gpu_per_trial: 1

The best hyperparameters for each experimental run are reported in Table S3.

Training employed the AdamW (69) optimizer with a linear learning rate scheduler and warmup over the first 10% of total steps. The total number of epochs was fixed at 100. Fixed hyperparameters included the scheduler type and warmup proportion, while the following were tuned jointly across the transformer and downstream components:

- **Transformer learning rate:** log-uniformly sampled from 1.0 × 10^-6^ to 1.0 × 10^-4^
- **Regressor learning rate:** log-uniformly sampled from 1.0 × 10^-5^ to 1.0 × 10^-3^
- **Transformer weight decay:** uniformly sampled from 0.0 to 0.1
- **Regressor weight decay:** uniformly sampled from 0.0 to 0.1
- **Transformer hidden dropout:** uniformly sampled from 0.1 to 0.5
- **Transformer attention dropout:** uniformly sampled from 0.1 to 0.5
- **Regressor dropout:** uniformly sampled from 0.1 to 0.5
- **Projector dropout:** uniformly sampled from 0.1 to 0.7

The performance of the full FlyUTR-RD model was compared to the spearman correlations of early mammalian models by downloading the analysis results table from Chu, et al (20) and finding the median value across the six spearman correlations for each tissue.

### Ribosome density data

RD measurements were obtained from ribosome profiling (Ribo-seq) and RNA-seq experiments conducted on *D. melanogaster* embryos at the maternal-to-zygotic transition (2–2.5 hours post-fertilization). Three biological replicates were used for both Ribo-seq and RNA-seq. Each replicate was normalized for gene length and library size. The normalized values for each datatype were averaged, and then a pseudo count of 1 was added to each gene. RD was calculated as the ratio of ribosome-protected fragment (RPF) density to mRNA abundance. Genes that did not have a raw count of at least one in all six datasets (3 replicates for both Ribo-seq and RNA-seq) were excluded from the analysis. Gene-level RD values were mapped to transcripts by identifying the highest expressed transcript for each gene using RNA-seq data from the same time point (39). Only transcripts with 5’ and 3’ UTR pairs that fit in the 512 token context window after tokenization were used for fine-tuning the model. For training, the RD values were double log transformed to force the model to differentiate between the bulk of the data rather than the outliers: log((log(RD+1)+1) (Figure S5).

### Model explainability analysis

We employed SHAP (SHapley Additive exPlanations) using the ferret(70) (v0.4.2) and shap(16) (v0.44.1) python packages to identify the sequence regions most influential for RD prediction. We first identified an appropriate token to use as the baseline by averaging all the token embeddings from the model (average embedding) and then isolating the token that was closest to this embedding in Euclidean space. We call this the ‘average token’. This approach balanced our desire for an average embedding with the desire to present an embedding seen by the model before. This mitigated the imbalanced SHAP values observed when the default [MASK] token was used.

SHAP values were calculated for each token position across all transcripts in the train, validation, and test datasets using the “partition” algorithm inside the Explainer and the ’average token’ as the mask_token in the TextMasker passed to the Explainer. We modified the Explainer class to properly handle the addition of the extra features before the MLP regression block (custom code found on our github).

### Global SHAP patterns across the 5’ and 3’ UTRs

For each transcript, we isolated the individual UTRs and interpolated their SHAP scores to 100 evenly spaced positions along each UTR using the interp1d function from *scipy* (*71*) (v1.15.3). This removed length differences, allowing direction comparison of SHAP distributions across transcripts. We then split the interpolated scores into positive and negative components, inserting NaN values in positions where scores of the opposite sign were removed. The absolute values of the negative SHAP scores were used so that SHAP magnitudes could be compared irrespective of direction. For each position along the UTRs, we calculated (i) the proportion of positive versus negative values and (ii) the average magnitudes of positive and negative SHAP scores.

Proportions were computed as the number of non-NaN values divided by the total number of transcripts. To compare SHAP magnitudes, we performed per-position statistical analyses using ordinary least squares (OLS) regression implemented with the *statsmodels* (*72*) library (v0.14.4). To increase statistical power, data from the train, validation, and test sets were combined for each UTR type. At each UTR position, we fit the model “value ∼ sign + dataset”, where *value* is the SHAP magnitude, *sign* indicates whether the score was positive or negative, and *dataset* specifies ‘train’, ‘val’, or ‘test’. P-values for *sign* were FDR corrected across positions.

Visualizations were generated using matplotlib (73) (v3.10.0). For each UTR, means were line-plotted against position for train, validation and test sets individually. Positions with significantly different SHAP magnitudes are marked in gold along the x-axis.

### Model-guided motif enrichment analysis

To perform the motif enrichment analysis, we first filtered out the sequences that the model poorly predicted, so these poor examples would not inject false hits into the motif enrichment analysis. We fit a LOWESS curve (statsmodels.nonparametric.lowess v0.14.4) to the residuals versus the experimental values and removed sequences where the residual was greater than two standard deviations away from the LOWESS curve.

Using the sequences that passed filtering, we identified all the binding locations for the RBPs within each transcript. Using apptainer (74) (v. 1.4.2-1.el9) to run the official docker image, we employed MAST(25) (v.5.5.7) from the MEME suite of tools using the Ray2013_rbp_Drosophila_melanogaster.meme motif collection(75) as a reference database (63 total motifs, 51 unique RBPs) and the --hit_list flag so that only the most probable nonoverlapping hits across all RBPs would remain. We also calculated the background nucleotide frequencies on the combined 5’ and 3’ UTRs in the training set and passed this into MAST with the -b flag.

We additionally used seaborn’s (v0.13.2) *clustermap* function to create a heatmap from the MAST motif correlations output from the MAST runs to visualize which RBP motifs are similar (Figure S3). We consider motifs with a correlation >= 0.8 as highly similar so we set the color scale max to 0.8 so that it was easier to identify all relationships above this threshold.

We then identified positive and negative high-importance regions in the transcripts. Using the distribution of SHAP scores from the training set, we call the high-importance regions in the train, validation, and test sets as those regions with scores greater than 1.25 standard deviations (train set derived) away from the train set mean. Positive high-importance regions (SHAP > train mean + 1.25 std) were compared to control regions (SHAP < train mean). Similarly, negative high-importance regions (SHAP < train mean – 1.25 std) were compared to control regions (SHAP > train mean).

To ensure fair comparison between the high-importance and control regions, we needed to make the length distributions between the sets comparable. We first order the lengths of the high-importance set so that the median comes first. We then alternate between one lower and one higher length so that the median number is first and the extreme values come last (‘ordered lengths list’). We then concatenate all the control sequences together and rechunk them by circling over the ‘ordered lengths list’, creating control regions that have the same length distribution as the high-importance set.

We then count the occurrences of each RBP for each sequence in the high-magnitude SHAP and control sets. For each RBP, we performed a Mann□Whitney U test (scipy v1.15.3) (71,76). P-values were corrected for multiple testing using the Benjamini□Hochberg method (77).

### Feature ablation and shuffling experiments

To evaluate the contributions of individual features to model performance, we conducted systematic ablation and shuffling analyses. All the experiments used the same dataset splits and evaluation metrics as the main model (MSE loss and Spearman correlation on the test set).

#### Feature ablation

In the full model, the input features comprised the concatenated 5’ and 3’ UTR sequences (processed by the transformer), GC content percentage, codon frequencies, lengths (computed separately for the 5’ UTR, coding sequence, and 3’ UTR), and mean free energy (MFE) for the entire transcript. For ablations excluding the sequences, we removed the transformer and retained only the MLP regressor. The MLP architecture consists of a linear layer (input dimension equal to the number of extra features, output dimension of 256), followed by ReLU activation, dropout, and a final linear layer (input dimension of 256, output dimension of 1).

The full unablated model is as follows:

- **FlyUTR-RD – Full model:** 5’ and 3’ UTR sequences + MFE + GC content + codon frequencies + lengths

The following ablations were performed:

1. **noMFE - No MFE:** 5’ and 3’ UTR sequences + GC content + codon frequencies + lengths
2. **noLengths - No lengths:** 5’ and 3’ UTR sequences + MFE + GC content + codon frequencies
3. **noGC - No GC content:** 5’ and 3’ UTR sequences + MFE + codon frequencies + lengths
4. **seq+Codons - No MFE, GC content, or lengths:** 5’ and 3’ UTR sequences + codon frequencies
5. **noCodons – No codon frequencies:** 5’ and 3’ UTR sequences + MFE + GC content + lengths
6. **seqOnly – No extra features:** 5’ and 3’ UTR sequences
7. **MLPall - No sequences (MLP):** MFE + GC content + codon frequencies + lengths
8. **MLPcodons - Codon frequencies only (MLP):** codon frequencies

An additional MLP ablation incorporated motif counts (from the MAST results) to approximate the sequence-derived motif information:

1. **MLPall+Motifs - No sequences + motif counts (MLP):** MFE + GC content + codon frequencies + lengths + motif counts

For transformer-based ablations (1–4), we used Ray Tune (67) with HyperOpt (68) to optimize all the same hyperparameters as the main model using the same Ray Tune parameters as the main model. For MLP-based ablations (5–7), we tuned

- **Learning rate:** log-uniformly sampled from 1.0 × 10^-5^ to 5.0 × 10^-3^
- **Weight decay:** uniformly sampled from 0.001 to 0.1
- **Dropout probability:** uniformly sampled from 0.1 to 0.9 using these ray tune parameters:

- ray_tune_samples: 300
- ray_tune_max_epochs: 100
- ray_tune_initial_points: 30
- ray_tune_grace_period: 5
- ray_tune_reduction_factor: 2
- ray_tune_cpu_per_trial: 1
- ray_tune_gpu_per_trial: 0

The best hyperparameters for each model are found in Table S3.

#### Feature shuffling

To assess the importance of features across transcripts, we performed shuffling experiments using the validation and test sets by recombining features from different transcripts while keeping others intact. Features were categorized into three groups: 5’ UTR sequences, 3’ UTR sequences, and non-sequence features (codon frequencies, MFE, GC content, and lengths). The following shuffles were applied using the FlyUTR-RD model:

1. Original 5’ UTR + original 3’ UTR + shuffled non-sequence features
2. Shuffled 5’ UTR + shuffled 3’ UTR + original non-sequence features
3. Shuffled 5’ UTR + original 3’ UTR + original non-sequence features
4. Original 5’ UTR + shuffled 3’ UTR + original non-sequence features

Here, "original" indicates features from the source transcript, while "shuffled" indicates features from another transcript in the same dataset split (validation or test). For each shuffled transcript, we computed the change in model prediction relative to the original prediction. Shuffles were performed using the best hyperparameters from the FlyUTR-RD model.

Because the number of tokens in the UTR affects token weighting (SHAP) in the transformer, shuffles 3 and 4 were restricted to pairs where the shuffled UTR token count was similar to the original. Specifically, we calculated the absolute token count difference across all candidate pairs and retained only those where the difference was less than one-third of the standard deviation of all differences.

### PAR-CLIP validation of Orb2

To test whether the model’s SHAP-derived Orb2 signal corresponded to experimental binding, we compared Orb2 motif predictions with Orb2B PAR-CLIP data from *Drosophila* S2 cells (GSE59611) (78). Orb2B is the most common protein isoform of the Orb2 gene whose motif was found to be enriched in the negative SHAP regions of both the 5’ and 3’ UTRs. Processed peak annotations were downloaded from CLIPdb (79) (http://clipdb.ncrnalab.org; bulk download: https://wj.qq.com/s2/10477617/4f84/v). Because Orb2 was identified in the negative SHAP category, we used low-SHAP windows from the 5′ and 3′ UTRs to define model-highlighted regions. Orb2 motif hits were obtained from MAST output across the train, validation, and test splits. PAR-CLIP peaks were restricted to UTR-assigned peaks, excluding CDS-overlapping sites, and genes were labeled as having 5′UTR, 3′UTR, or any-UTR Orb2B binding.

For UTR-level validation, genes were classified into three mutually exclusive groups: Group A, no Orb2 MAST hit; Group B, Orb2 MAST hit without overlap with a low-SHAP window; and Group C, Orb2 MAST hit overlapping a low-SHAP window. Grouping was performed for any UTR, 5′UTR only, and 3′UTR only, with isoform-level assignments collapsed to gene level using the priority C > B > A. The primary test of SHAP added value compared Group C with Group B, asking whether SHAP-selected motif-containing genes were more likely than motif-only genes to be supported by PAR-CLIP. Enrichment was assessed using one-sided Fisher’s exact tests, and odds ratios, precision, recall, and contingency tables were recorded.

For position-level validation, transcript-local Orb2 MAST hit coordinates were converted to genomic intervals using UTR exon structures from the same Ensembl GTF used to isolate the UTR sequences from the genome. Motif sites were classified as inside or outside low-SHAP windows, collapsed to unique genomic intervals, and scored for overlap with Orb2B PAR-CLIP peaks on the same chromosome and strand. Position-level enrichment was then tested by comparing motif sites inside versus outside SHAP windows for all UTRs combined, 5′ UTRs only, and 3′ UTRs only using one-sided Fisher’s exact tests.

### Naïve motif enrichment analysis

To derive balanced high- and low-RD subsets for downstream analyses, transcripts were sorted in ascending order by RD rate. The sorted dataset was partitioned into five quantiles. The two lowest quantiles (lowest 40% of rates) constituted the low-RD group, while the two highest quantiles (highest 40% of rates) formed the high-RD group; the central quantile was discarded to ensure group separation.

As in the model-guided analysis (above), MAST was used with the --hit-list flag to identify the best nonoverlapping RBPs bound to each transcript. To mitigate length biases, UTRs were segmented into nonoverlapping pseudofragments of ∼25 nucleotides (median length). Specifically, breakpoints were searched for every 25 nucleotides and placed at the nearest non-motif-hit position to ensure that fragmentation did not disrupt a recognized motif.

For the 5’ and 3’ UTRs separately, high- and low-RD segment-level RBP counts were compared using nonparametric Mann□Whitney U tests (Scipy v1.13.1). Benjamini□Hochberg false discovery rate (FDR) correction was applied across the RBPs using Statsmodels v0.14.4.

### Linear model for motif association analysis

To provide an interpretable baseline for motif-level effects, we trained a linear regression model using ordinary least squares (scikit-learn) to predict ribosome density (RD) from transcript-level features. Input features included motif occurrence counts per transcript (one feature per RNA-binding protein motif) together with the additional sequence-derived covariates included in FlyUTR-RD (e.g. MFE with 5’, 3’, and CDS specific length and GC content features).

The model was trained and evaluated on the training dataset only, with predictions generated on the same data, as the objective was not predictive performance but estimation of motif-associated effects. Motifs were ranked according to their t-statistics and direction of effect as determined by the sign of the corresponding regression coefficient. These motif rankings were used for comparison with those derived from naïve enrichment analysis and SHAP-based attribution methods.

### Decay (half-life) analysis

Using the same methodology described in our previous publication (39), we extended our decay rate modelling from 263 zygotic transcripts to 655 transcripts with high quality results. Of the 655 transcripts, 615 had UTR pairs that could fit into the 512 token context window of the model after tokenization. Using our same extended pretraining model described above (GENA LM Fly UTR), we then fine-tuned on this small set of mRNA decay rates. We first converted the decay rates to half-life:

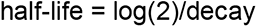

and then log transformed them. We then ran the same pipeline built for the RD analysis with only a few variations:

1. We did not need to map the rates from genes to transcripts because the data are already transcript specific.
2. During ray tune hyperparameter optimization (Table S3), we changed the optimization objective to minimizing the validation loss rather than maximizing the Spearman coefficient as this provided better hyperparameters. The training objective remained unchanged at minimizing the loss.
3. We selected one of the ablation models (noCodons) for the downstream motif enrichment analyses rather than the full feature set genomic foundation model.

The best hyperparameters for each experimental run are reported in Table S4.

## Statistical analysis

All statistical analyses were performed in Python (version 3.10.16). Multiple testing correction was applied using the Benjamini□Hochberg method. Model performance was evaluated using Spearman correlation coefficients, coefficients of determination, and mean squared error (MSE) metrics.

## Data availability

The raw RNA-seq data used to produce the decay estimates are publicly available at ArrayExpress under accession number: E-MTAB-11580 (https://www.ebi.ac.uk/biostudies/ArrayExpress/studies/E-MTAB-11580). The decay estimates themselves and the RD data are available via Zenodo at: https://doi.org/10.5281/zenodo.20446580 and are also included in the github repository: https://github.com/ManchesterBioinference/mRNA_LLM. We downloaded the *Drosophila melanogaster* reference genome from Ensembl (https://ftp.ensembl.org/pub/rapid-release/species/Drosophila_melanogaster/GCA_000001215.4/flybase/genome/Drosophila_ melanogaster-GCA_000001215.4-unmasked.fa.gz). The transcript annotations were downloaded from Ensembl (https://ftp.ensembl.org/pub/rapid-release/species/Drosophila_melanogaster/GCA_000001215.4/flybase/geneset/2022_07/Dro sophila_melanogaster-GCA_000001215.4-2022_07-genes.gtf.gz).

The ortholog data were downloaded from DrosOMA (https://drosoma.unil.ch/All/oma-groups.txt.gz). The file mapping OMA ids to RefSeq accessions was downloaded from the DrosOMA website (https://drosoma.unil.ch/All/oma-ncbi.txt.gz). The gene2refseq file that maps gene symbols and RefSeq accessions was downloaded from NCBI (https://ftp.ncbi.nlm.nih.gov/gene/DATA/gene2refseq.gz). The file mapping gene symbols and Flybase IDs was downloaded from flybase by passing the unique gene symbols from the gene2refseq file to the Flybase ID Validator tool (https://flybase.org/convert/id).

The Ray RBP motif database is included in the docker image (https://hub.docker.com/r/memesuite/memesuite) under this path: /opt/meme/share/meme-5.5.7/db/motif_databases/RNA/Ray2013_rbp_Drosophila_melanogaster.meme

The GENA-LM fly model is available on huggingface (https://huggingface.co/AIRI-Institute/gena-lm-bert-base-fly).

The source data for the results of the RD benchmarking analysis performed on six mammalian models in three different tissues can be downloaded from the article in Nature Machine Intelligence (https://static-content.springer.com/esm/art%3A10.1038%2Fs42256-024-00823-9/MediaObjects/42256_2024_823_MOESM3_ESM.xlsx)

## Code availability

All of the analyses were incorporated into a reproducible pipeline using the Data Version Control (DVC v3.63.0) tool and is available on our github at https://github.com/ManchesterBioinference/mRNA_LLM and via Zenodo at: https://doi.org/10.5281/zenodo.20446580. All necessary data is either included in the github repo or is programmatically downloaded from publicly available databases by the pipeline. The RD analysis can be found under the “translationEfficiency” branch, and the half-life (decay) analysis can be found under the “decay” branch.

MAST, from the MEME suite of tools, was accessed via apptainer(74) using MEME’s official docker image (https://hub.docker.com/r/memesuite/memesuite): docker://memesuite/memesuite:5.5.7.

## Supporting information

2h_ribosome_density_data

decay_estimates

## Acknowledgements

Funding is a Wellcome Trust Discovery Award to HLA and MR (227415/Z/23/Z) and a BBSRC grant to HLA and MPA (BB/X007294/1). Various LLMs assisted in code creation and manuscript editing.

## Author information

### Authors and affiliations

#### Contributions

L.B. analysed the data, built the processing pipeline and wrote the manuscript. D.C. generated the experimental RD data. Y.S. calculated the mRNA decay estimates from the experimental decay data and reviewed the manuscript. D.C., M.A. and H.L.A. contributed biological insights. D.C., M.A., H.L.A. and M.R. edited the manuscript. H.L.A. and M.R. organized and supervised the project.

#### Corresponding authors

Correspondence to Hilary Ashe or Magnus Rattray

## Ethics declarations

### Ethics approval

Not applicable.

### Consent for publication

Not applicable.

## Supplementary figures

**Figure S1.**
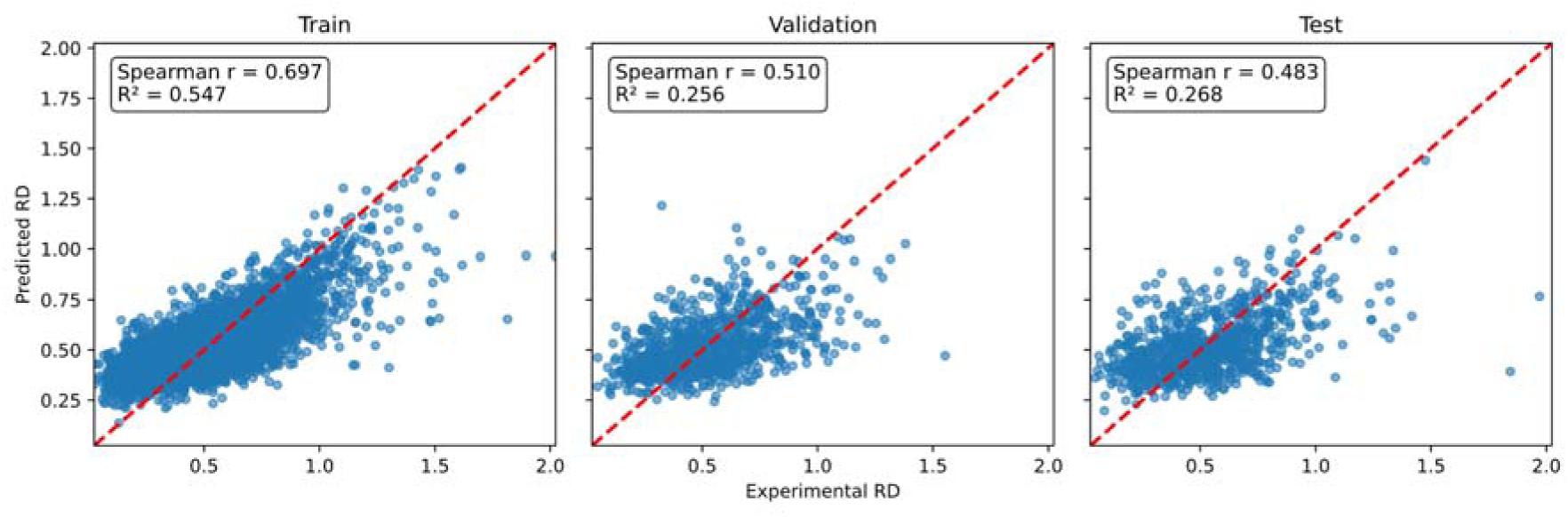
Scatter plots of the FlyUTR-RD model’s predictions versus the experimental RD values for the train, validation, and test sets.

**Figure S2.**
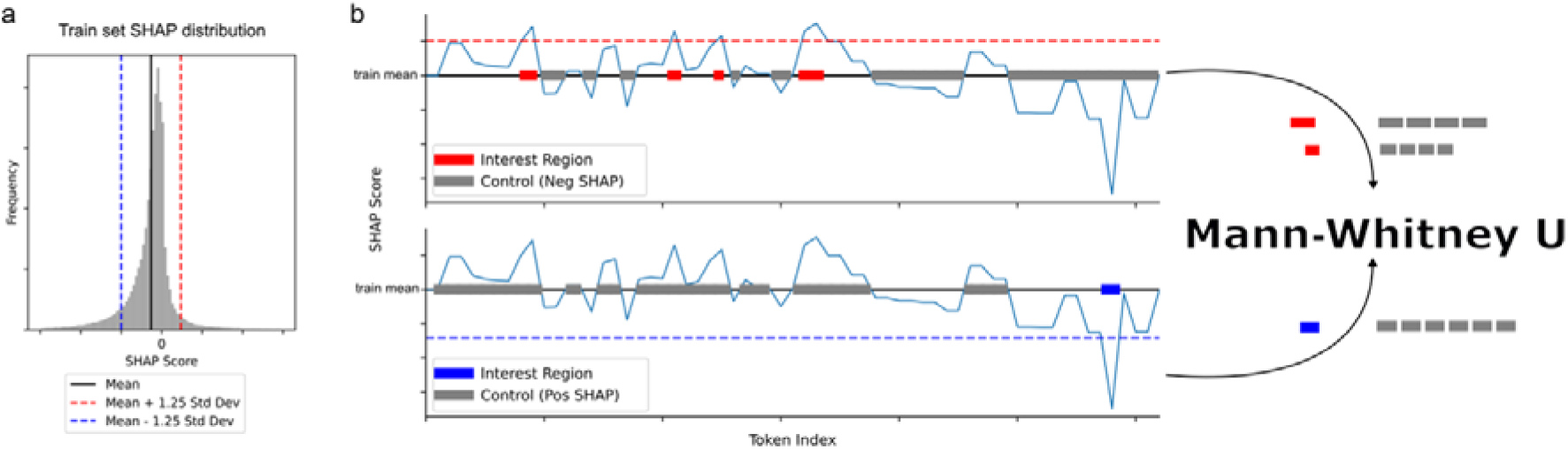
Schematic for identifying high-importance SHAP regions. a) Thresholds are identified from the train set SHAP distribution. b) The transcript body is visualized on the training set mean. We call high-importance regions in the train, validation, and test sets as those regions with scores greater than 1.25 standard deviations (train set derived) away from the train set mean. Positive high-importance regions (SHAP > train mean + 1.25 std; above the red dashed lines) are compared to control regions (SHAP < train mean). Similarly, negative high-importance regions (SHAP < train mean – 1.25 std; below the blue dashed lines) are compared to control regions (SHAP > train mean).

**Figure S3.**
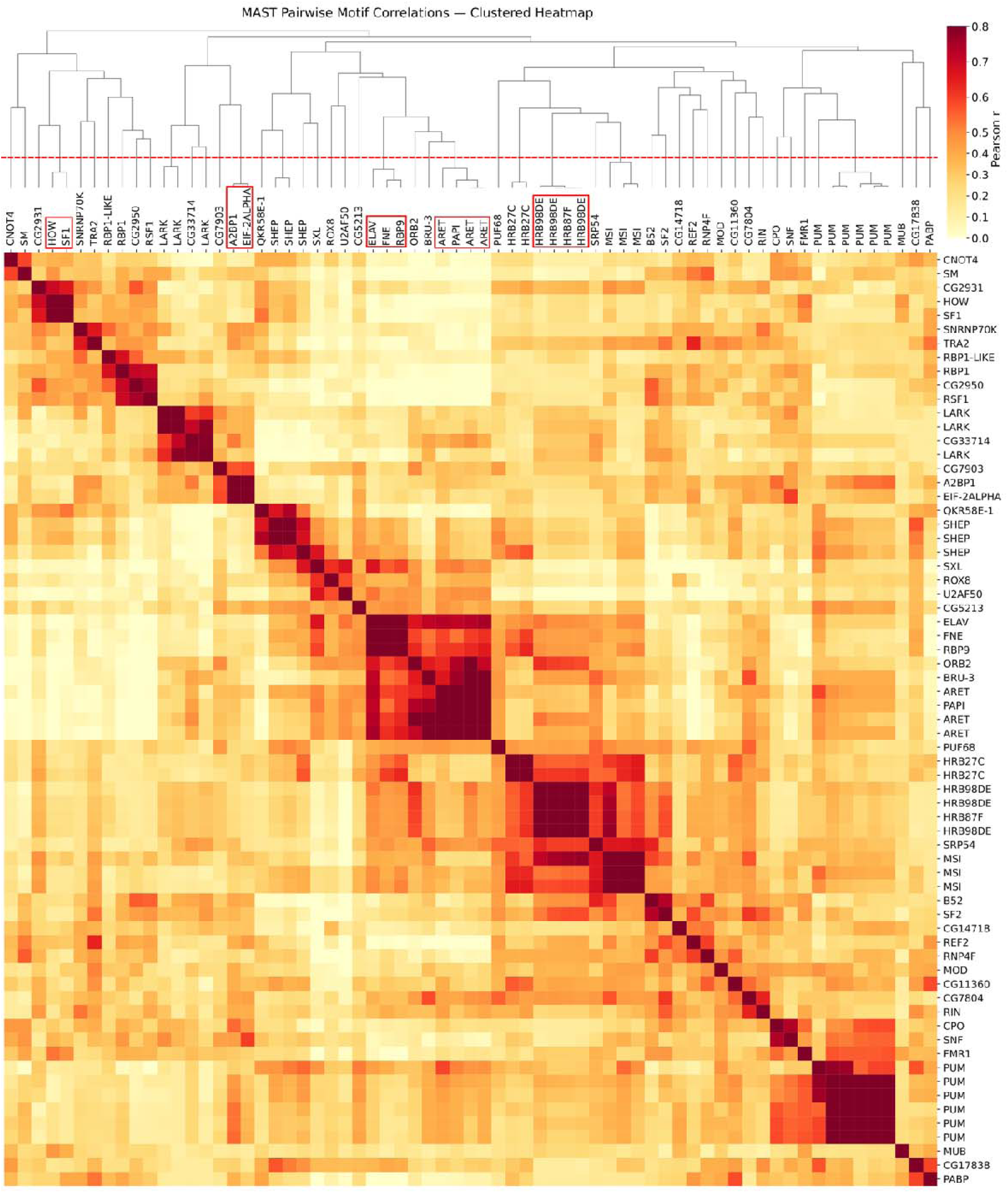
Heatmap of the MAST motif correlations. The branches below the red dashed line intersecting the dendrogram indicate a >= 80% Pearson correlation between the RBP motifs. Groups below this line that have more than one unique RBP are boxed in red to highlight RBPs with similar motifs.

**Figure S4.**
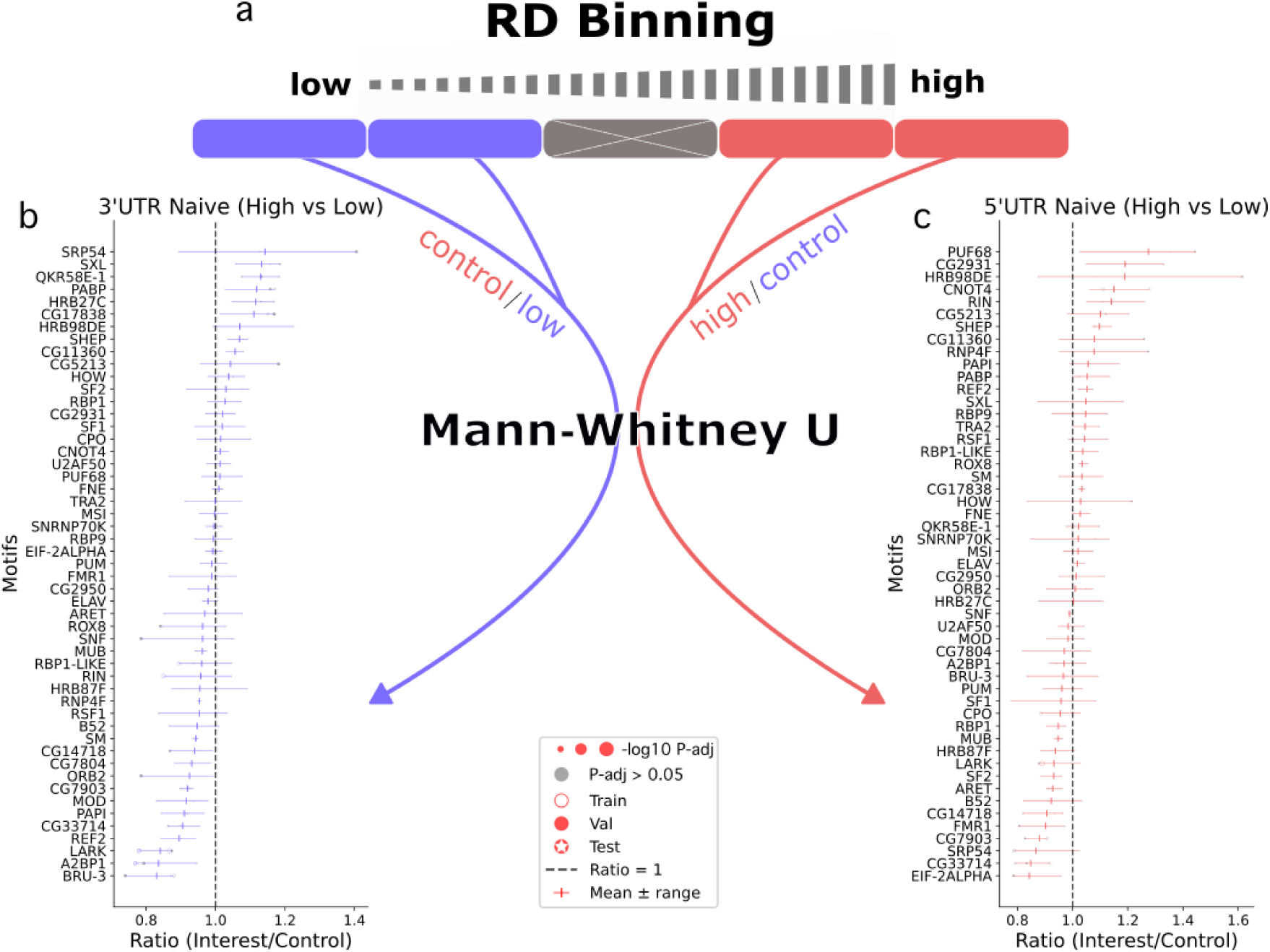
Naïve high versus low RD analysis. a) Schematic of the transcript binning used in the ‘naïve’ high- versus low-RD AME analysis. b, c) Mann L Whitney U enrichment results for the low (b) and high (c) RD transcripts.

**Figure S5.**
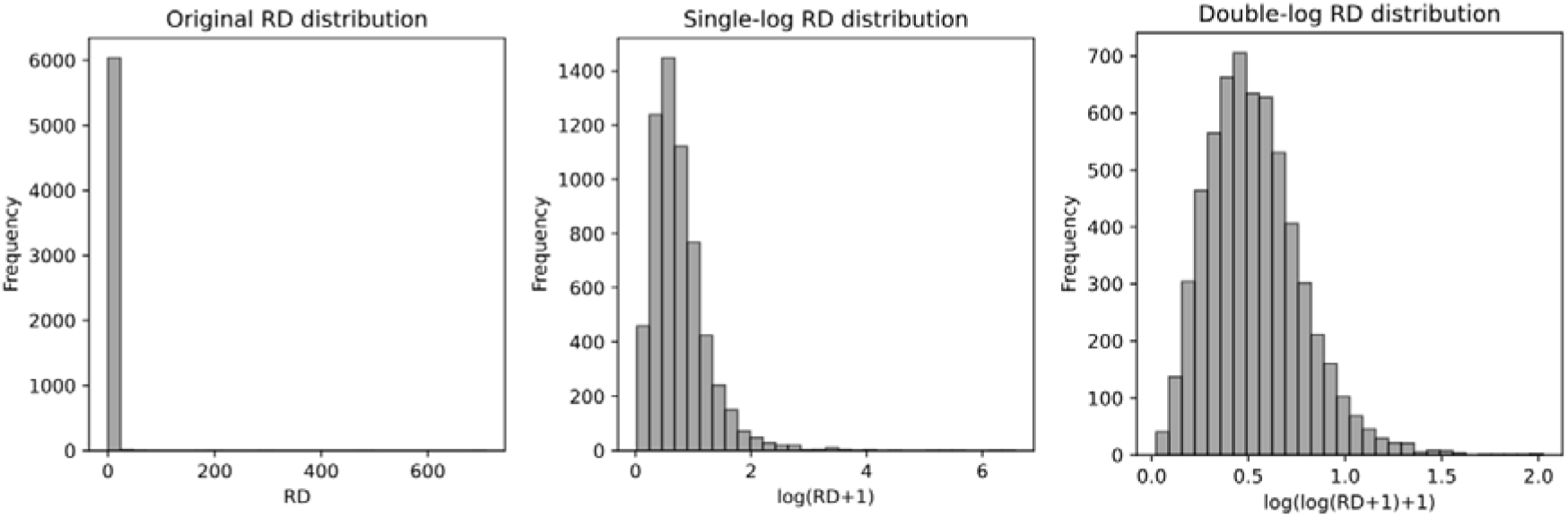
Ribosome density score distributions before and after transformation.

## Supplementary tables

**Table S1.**
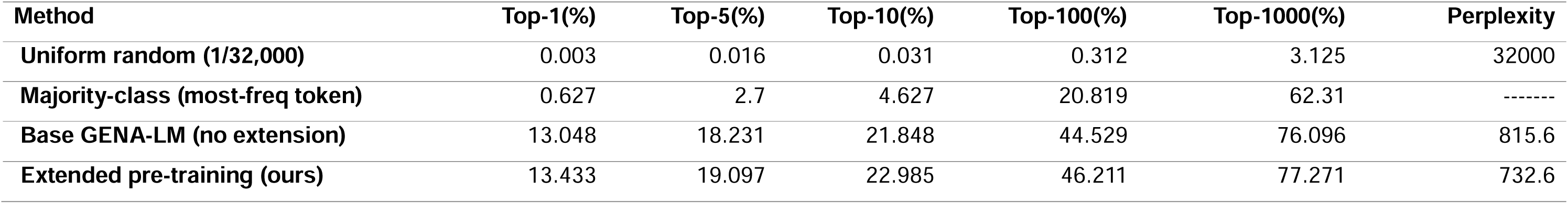
Token prediction accuracy. The percentage of predictions where the true token is in the Top-1, Top-5, Top-10, Top-100, or Top-1000 of predicted tokens. Comparison between different token selection methods, uniformly random, most frequent token, base GENA-LM Fly (no extension), or GENA-LM Fly UTR (extended pertaining; ours).

**Table S2.** Model Metrics.

| Model | test R<sup>2</sup> | test Spearman<br>$\rho$ | train MSE | val MSE | test MSE |
| --- | --- | --- | --- | --- | --- |
| FlyUTR-RD | 0.2681 | 0.4828 | 0.039323 | 0.040817 | 0.043008 |
| noCodons* | 0.0343 | 0.2210 | 0.062704 | 0.050262 | 0.057079 |
| noGC | 0.2416 | 0.4557 | 0.035233 | 0.04049 | 0.044536 |
| noLengths | 0.2096 | 0.4490 | 0.043476 | 0.042449 | 0.046434 |
| noMFE | 0.2493 | 0.4798 | 0.039868 | 0.040003 | 0.044226 |
| seq+Codons | 0.1226 | 0.4640 | 0.045631 | 0.039756 | 0.051465 |
| seqOnly* | -0.1268 | 0.1931 | 0.05964 | 0.050206 | 0.066627 |
| MLPcodons | 0.2205 | 0.4404 | 0.05636 | 0.043396 | 0.045658 |
| MLPall | 0.2510 | 0.4626 | 0.051047 | 0.043078 | 0.045177 |
| MLPall+Motifs | 0.2303 | 0.4526 | 0.054611 | 0.044521 | 0.05042 |

Coefficient of determination (R<sup>2</sup>) and Spearman correlation coefficient ( $\rho$ ) for the test set and the MSE for the train, validation (val), and test sets for each model.
\* these model started to overfit quickly, so early stopping terminated the training sooner than the other models, preventing learning.

**Table S3.** Best RD hyperparameters identified by Ray Tune with HyperOpt for each model.

|  | bert_lr | classifier_lr | bert_weight_decay | classifier_weight_decay | bert_hidden_dropout | bert_atten_dropout | classifier_dropout | projector_dropout |
| --- | --- | --- | --- | --- | --- | --- | --- | --- |
| FlyUTR-RD | 8.684526659564706e-05 | 0.000992716507960959 | 0.05945982266288608 | 0.06953767497101772 | 0.21995700110481797 | 0.43374654573885274 | 0.4228761858931757 | 0.10533686891488281 |
| noLengths | 4.0324863501064064e-05 | 0.0009948618156170358 | 0.07121862526176946 | 0.06512150673036911 | 0.3060516079997689 | 0.2375750410259943 | 0.23707813988731752 | 0.39211037828986717 |
| seq+Codons | 3.049793528732112e-05 | 0.00038843962179367647 | 0.0578161813106169 | 0.08847081039151136 | 0.37887155309561477 | 0.19473410742876213 | 0.17751588706532484 | 0.400967693124663 |
| seqOnly | 7.184337918860248e-05 | 2.4120455810509182e-05 | 0.028422139272116172 | 0.005268440733890316 | 0.1283785833154604 | 0.39495198739783444 | 0.30162902314338724 | 0.679509034281437 |
| noMFE | 7.677032374882764e-06 | 0.000977056745878954 | 0.09055158652107101 | 0.06765989035324677 | 0.27036326183512716 | 0.27493868511300495 | 0.1382787799834731 | 0.3145456709703972 |
| noCodons | 5.5026940980111536e-05 | 1.007488301598011e-05 | 0.09950280245155382 | 0.07102495316730657 | 0.12364444141121064 | 0.21289446531364953 | 0.47119707623344165 | 0.6827732103421423 |
| noGC | 5.206327059333067e-06 | 0.000983314294722826 | 0.001983044773115374 | 0.05521254188737569 | 0.4362881220855709 | 0.3259015078171584 | 0.19654481823660083 | 0.5589147335429167 |
| MLPall+Motifs |  | 0.0038491183337653534 |  | 0.05853100581866877 |  |  | 0.7406828295461403 |  |
| MLPcodons |  | 0.003288690933361583 |  | 0.019210395423453056 |  |  | 0.7784108919655274 |  |
| MLPall |  | 0.003030338952388465 |  | 0.08311893020380384 |  |  | 0.7520468593720935 |  |

**Table S4.**
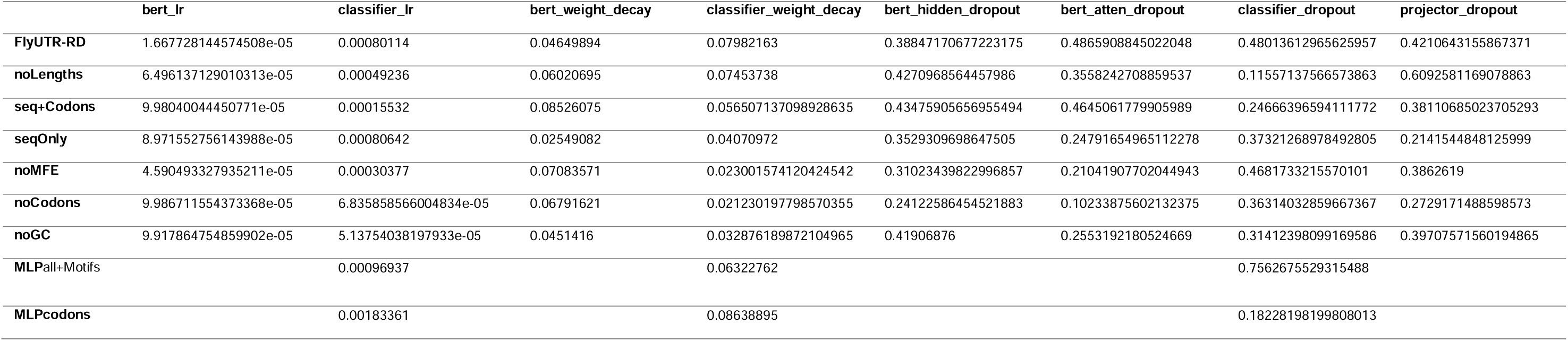
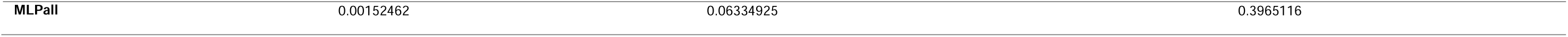
Best half-life hyperparameters identified by Ray Tune with HyperOpt for each model.

## References

1. Harrison MM, Marsh AJ, Rushlow CA. Setting the stage for development: the maternal-to-zygotic transition in Drosophila. Genetics. 2023 Oct 1;225(2):iyad142. doi:10.1093/genetics/iyad142

2. Teixeira FK, Lehmann R. Translational Control during Developmental Transitions. Cold Spring Harb Perspect Biol. 2019 Jun;11(6):a032987. doi:10.1101/cshperspect.a032987 PubMed PMID: 30082467; PubMed Central PMCID: PMC6546043.

3. Kronja I, Yuan B, Eichhorn S, Dzeyk K, Krijgsveld J, Bartel DP, et al. Widespread changes in the posttranscriptional landscape at the Drosophila oocyte-to-embryo transition. Cell Rep. 2014 Jun 12;7(5):1495–508. doi:10.1016/j.celrep.2014.05.002 PubMed PMID: 24882012; PubMed Central PMCID: PMC4143395.

4. Benoit B, He CH, Zhang F, Votruba SM, Tadros W, Westwood JT, et al. An essential role for the RNA-binding protein Smaug during the Drosophila maternal-to-zygotic transition. Development. 2009 Mar 15;136(6):923–32. doi:10.1242/dev.031815 PubMed PMID: 19234062; PubMed Central PMCID: PMC2727558.

5. Rodriguez-Martinez A, Young-Baird SK. Polysome profiling is an extensible tool for the analysis of bulk protein synthesis, ribosome biogenesis, and the specific steps in translation. Mol Biol Cell. 2025 Mar 14;36(4):mr2. doi:10.1091/mbc.E24-08-0341 PubMed PMID: 40042939; PubMed Central PMCID: PMC12005114.

6. Ichinose T, Kondo S, Kanno M, Shichino Y, Mito M, Iwasaki S, et al. Translational regulation enhances distinction of cell types in the nervous system. eLife. 2024 Jun 13;12. doi:10.7554/eLife.90713.2

7. Chen X, Dickman D. Development of a tissue-specific ribosome profiling approach in Drosophila enables genome-wide evaluation of translational adaptations. PLOS Genetics. 2017 Dec 1;13(12):e1007117. doi:10.1371/journal.pgen.1007117

8. Schmidt EK, Clavarino G, Ceppi M, Pierre P. SUnSET, a nonradioactive method to monitor protein synthesis. Nat Methods. 2009 Apr;6(4):275–7. doi:10.1038/nmeth.1314 PubMed PMID: 19305406.

9. Zheng D, Persyn L, Wang J, Liu Y, Ulloa-Montoya F, Cenik C, et al. Predicting the translation efficiency of messenger RNA in mammalian cells. Nat Biotechnol. 2025 Jul 25;1–14. doi:10.1038/s41587-025-02712-x

10. Agarwal V, Kelley DR. The genetic and biochemical determinants of mRNA degradation rates in mammals. Genome Biology. 2022 Nov 23;23(1):245. doi:10.1186/s13059-022-02811-x

11. Yang Y, Li G, Pang K, Cao W, Zhang Z, Li X. Deciphering 3’UTR Mediated Gene Regulation Using Interpretable Deep Representation Learning. Advanced Science. n/a(n/a):2407013. doi:10.1002/advs.202407013

12. Fishman V, Kuratov Y, Shmelev A, Petrov M, Penzar D, Shepelin D, et al. GENA-LM: a family of open-source foundational DNA language models for long sequences. Nucleic Acids Res. 2025 Jan 27;53(2):gkae1310. doi:10.1093/nar/gkae1310

13. Dalla-Torre H, Gonzalez L, Mendoza-Revilla J, Carranza NL, Grzywaczewski AH, Oteri F, et al. The Nucleotide Transformer: Building and Evaluating Robust Foundation Models for Human Genomics [Internet]. bioRxiv; 2023 [cited 2024 Oct 1]. p. 2023.01.11.523679. Available from: https://www.biorxiv.org/content/10.1101/2023.01.11.523679v3 doi:10.1101/2023.01.11.523679

14. Nguyen E, Poli M, Faizi M, Thomas A, Birch-Sykes C, Wornow M, et al. HyenaDNA: Long-Range Genomic Sequence Modeling at Single Nucleotide Resolution [Internet]. arXiv; 2023 [cited 2025 Mar 11]. Available from: http://arxiv.org/abs/2306.15794 doi:10.48550/arXiv.2306.15794

15. Jung AJ, Zhu H, Gao AJ, Li R, Slobodyanyuk M, Chu V, et al. FlashRNA: An Efficient Model for Regulatory Genomics [Internet]. bioRxiv; 2025 [cited 2025 Oct 21]. p. 2025.10.14.682350. Available from: https://www.biorxiv.org/content/10.1101/2025.10.14.682350v1 doi:10.1101/2025.10.14.682350

16. Lundberg S, Lee SI. A Unified Approach to Interpreting Model Predictions [Internet]. arXiv; 2017 [cited 2025 Nov 21]. Available from: http://arxiv.org/abs/1705.07874 doi:10.48550/arXiv.1705.07874

17. Simonyan K, Vedaldi A, Zisserman A. Deep Inside Convolutional Networks: Visualising Image Classification Models and Saliency Maps [Internet]. arXiv; 2014 [cited 2025 Nov 13]. Available from: http://arxiv.org/abs/1312.6034 doi:10.48550/arXiv.1312.6034

18. Sundararajan M, Taly A, Yan Q. Axiomatic Attribution for Deep Networks. In: Proceedings of the 34th International Conference on Machine Learning [Internet]. PMLR; 2017 [cited 2025 Nov 13]. p. 3319–28. Available from: https://proceedings.mlr.press/v70/sundararajan17a.html

19. Ribeiro MT, Singh S, Guestrin C. ‘Why Should I Trust You?’: Explaining the Predictions of Any Classifier. In: Proceedings of the 22nd ACM SIGKDD International Conference on Knowledge Discovery and Data Mining [Internet]. New York, NY, USA: Association for Computing Machinery; 2016 [cited 2025 Nov 13]. p. 1135–44. (KDD ’16). Available from: https://dl.acm.org/doi/10.1145/2939672.2939778 doi:10.1145/2939672.2939778

20. Chu Y, Yu D, Li Y, Huang K, Shen Y, Cong L, et al. A 5’ UTR Language Model for Decoding Untranslated Regions of mRNA and Function Predictions. Nat Mach Intell. 2024 Apr;6(4):449–60. doi:10.1038/s42256-024-00823-9 PubMed PMID: 38855263; PubMed Central PMCID: PMC11155392.

21. Sharp PM, Li WH. An evolutionary perspective on synonymous codon usage in unicellular organisms. J Mol Evol. 1986 Dec 1;24(1):28–38. doi:10.1007/BF02099948

22. Gingold H, Pilpel Y. Determinants of translation efficiency and accuracy. Mol Syst Biol. 2011 Apr 12;7:481. doi:10.1038/msb.2011.14 PubMed PMID: 21487400; PubMed Central PMCID: PMC3101949.

23. Jacobson A. 16 Poly(A) Metabolism and Translation: The Closed-loop Model. Cold Spring Harbor Monograph Archive. 1996 Jan 1;30(0):451–80. doi:10.1101/0.451-480

24. Romagnoli A, D’Agostino M, Ardiccioni C, Maracci C, Motta S, La Teana A, et al. Control of the eIF4E activity: structural insights and pharmacological implications. Cell Mol Life Sci. 2021 Sep 19;78(21–22):6869–85. doi:10.1007/s00018-021-03938-z PubMed PMID: 34541613; PubMed Central PMCID: PMC8558276.

25. Bailey TL, Gribskov M. Combining evidence using p-values: application to sequence homology searches. Bioinformatics. 1998;14(1):48–54. doi:10.1093/bioinformatics/14.1.48 PubMed PMID: 9520501.

26. Hernández G, Miron M, Han H, Liu N, Magescas J, Tettweiler G, et al. Mextli Is a Novel Eukaryotic Translation Initiation Factor 4E-Binding Protein That Promotes Translation in Drosophila melanogaster. Mol Cell Biol. 2013 Aug;33(15):2854–64. doi:10.1128/MCB.01354-12 PubMed PMID: 23716590; PubMed Central PMCID: PMC3719689.

27. Ghosh S, Thomas SE, Abraham LM, Vaughn JC. Regulation of Expression for the RNP-4F Splicing Assembly Factor in the Fruit-Fly Drosophila melanogaster. Open Journal of Animal Sciences. 2015 Oct 9;5(4):418–28. doi:10.4236/ojas.2015.54044

28. Sanford JR, Gray NK, Beckmann K, Cáceres JF. A novel role for shuttling SR proteins in mRNA translation. Genes Dev. 2004 Apr 1;18(7):755–68. doi:10.1101/gad.286404 PubMed PMID: 15082528; PubMed Central PMCID: PMC387416.

29. Maslon MM, Heras SR, Bellora N, Eyras E, Cáceres JF. The translational landscape of the splicing factor SRSF1 and its role in mitosis. Nahum S, editor. eLife. 2014 May 6;3:e02028. doi:10.7554/eLife.02028

30. Carreira-Rosario A, Bhargava V, Hillebrand J, Kollipara RK, Ramaswami M, Buszczak M. Repression of Pumilio Protein Expression by Rbfox1 Promotes Germ Cell Differentiation. Dev Cell. 2016 Mar 7;36(5):562–71. doi:10.1016/j.devcel.2016.02.010 PubMed PMID: 26954550; PubMed Central PMCID: PMC4785839.

31. Lie YS, Macdonald PM. Apontic binds the translational repressor Bruno and is implicated in regulation of oskar mRNA translation. Development. 1999 Mar 15;126(6):1129–38. doi:10.1242/dev.126.6.1129

32. Delaunay J, Le Mée G, Ezzeddine N, Labesse G, Terzian C, Capri M, et al. The Drosophila Bruno paralogue Bru-3 specifically binds the EDEN translational repression element. Nucleic Acids Res. 2004;32(10):3070–82. doi:10.1093/nar/gkh627 PubMed PMID: 15181172; PubMed Central PMCID: PMC434433.

33. Khan M ‘Repon’, Li L, Pérez-Sánchez C, Saraf A, Florens L, Slaughter BD, et al. Amyloidogenic oligomerization transforms Drosophila Orb2 from a translation repressor to an activator. Cell. 2015 Dec 3;163(6):1468–83. doi:10.1016/j.cell.2015.11.020 PubMed PMID: 26638074; PubMed Central PMCID: PMC4674814.

34. Goldstrohm AC, Tanaka Hall TM, McKenney KM. Post-transcriptional Regulatory Functions of Mammalian Pumilio Proteins. Trends Genet. 2018 Dec;34(12):972–90. doi:10.1016/j.tig.2018.09.006 PubMed PMID: 30316580; PubMed Central PMCID: PMC6251728.

35. Grzejda D, Hess A, Rezansoff A, Gorey S, Carrasco J, Alfonso-Gonzalez C, et al. Pumilio differentially binds to mRNA 3′ UTR isoforms to regulate localization of synaptic proteins. EMBO reports. 2025 Apr 7;26(7):1792–815. doi:10.1038/s44319-025-00401-z

36. Kim-Ha J, Kim J, Kim YJ. Requirement of RBP9, a Drosophila Hu homolog, for regulation of cystocyte differentiation and oocyte determination during oogenesis. Mol Cell Biol. 1999 Apr;19(4):2505–14. doi:10.1128/MCB.19.4.2505 PubMed PMID: 10082516; PubMed Central PMCID: PMC84043.

37. Buddika K, Ariyapala IS, Haugan MA, Riffert D, Sokol NS. Canonical nucleators are dispensable for stress granule assembly in Drosophila intestinal progenitors. J Cell Sci. 2020 May 18;133(10):jcs243451. doi:10.1242/jcs.243451 PubMed PMID: 32265270; PubMed Central PMCID: PMC7325430.

38. Penalva LOF, Sánchez L. RNA Binding Protein Sex-Lethal (Sxl) and Control of Drosophila Sex Determination and Dosage Compensation. Microbiology and Molecular Biology Reviews. 2003 Sep;67(3):343–59. doi:10.1128/mmbr.67.3.343-359.2003

39. Beadle LF, Love JC, Shapovalova Y, Artemev A, Rattray M, Ashe HL. Combined modelling of mRNA decay dynamics and single-molecule imaging in the Drosophila embryo uncovers a role for P-bodies in 5′ to 3′ degradation. PLOS Biology. 2023 Jan 17;21(1):e3001956. doi:10.1371/journal.pbio.3001956

40. Samuels TJ, Torley EJ, Nadmitova V, Naden EL, Blair PE, Hernandez Frometa FA, et al. Destabilisation of bam transcripts terminates the mitotic phase of Drosophila female germline differentiation. Development. 2025 Mar 11;152(5):DEV204405. doi:10.1242/dev.204405

41. Guo X, Sun Y, Azad T, Janse van Rensburg HJ, Luo J, Yang S, et al. Rox8 promotes microRNA-dependent yki messenger RNA decay. Proceedings of the National Academy of Sciences. 2020 Dec;117(48):30520–30. doi:10.1073/pnas.2013449117

42. Giuliani G, Giuliani F, Volk T, Rabouille C. The Drosophila RNA-binding protein HOW controls the stability of dgrasp mRNA in the follicular epithelium. Nucleic Acids Research. 2014 Feb;42(3):1970–86. doi:10.1093/nar/gkt1118

43. McDermott SM, Meignin C, Rappsilber J, Davis I. Drosophila Syncrip binds the gurken mRNA localisation signal and regulates localised transcripts during axis specification. Biol Open. 2012 Apr 11;1(5):488–97. doi:10.1242/bio.2012885 PubMed PMID: 23213441; PubMed Central PMCID: PMC3507208.

44. Sofola O, Sundram V, Ng F, Kleyner Y, Morales J, Botas J, et al. The Drosophila FMRP and LARK RNA-Binding Proteins Function Together to Regulate Eye Development and Circadian Behavior. J Neurosci. 2008 Oct 8;28(41):10200–5. doi:10.1523/JNEUROSCI.2786-08.2008 PubMed PMID: 18842880.

45. Grams R, Korge G. The *mub* gene encodes a protein containing three KH domains and is expressed in the mushroom bodies of *Drosophila melanogaster*. Gene. 1998 Jul 17;215(1):191–201. doi:10.1016/S0378-1119(98)00251-0

46. Kini HK, Vishnu MR, Liebhaber SA. Too much PABP, too little translation. J Clin Invest. 2010 Sep 1;120(9):3090–3. doi:10.1172/JCI44091 PubMed PMID: 20739750; PubMed Central PMCID: PMC2929741.

47. Gu S, Jeon HM, Nam SW, Hong KY, Rahman MS, Lee JB, et al. The flip-flop configuration of the PABP-dimer leads to switching of the translation function. Nucleic Acids Res. 2021 Dec 14;50(1):306–21. doi:10.1093/nar/gkab1205 PubMed PMID: 34904669; PubMed Central PMCID: PMC8754640.

48. Cline TW, Rudner DZ, Barbash DA, Bell M, Vutien R. Functioning of the Drosophila integral U1/U2 protein Snf independent of U1 and U2 small nuclear ribonucleoprotein particles is revealed by snf+ gene dose effects. Proc Natl Acad Sci U S A. 1999 Dec 7;96(25):14451–8. doi:10.1073/pnas.96.25.14451 PubMed PMID: 10588726; PubMed Central PMCID: PMC24457.

49. Blanchette M, Labourier E, Green RE, Brenner SE, Rio DC. Genome-Wide Analysis Reveals an Unexpected Function for the *Drosophila* Splicing Factor U2AF50 in the Nuclear Export of Intronless mRNAs. Molecular Cell. 2004 Jun 18;14(6):775–86. doi:10.1016/j.molcel.2004.06.012

50. Liu L, Qi H, Wang J, Lin H. PAPI, a novel TUDOR-domain protein, complexes with AGO3, ME31B and TRAL in the nuage to silence transposition. Development. 2011 May;138(9):1863–73. doi:10.1242/dev.059287 PubMed PMID: 21447556; PubMed Central PMCID: PMC3074456.

51. Carrasco J, Mateos F, Hilgers V. A critical developmental window for ELAV/Hu-dependent mRNA signatures at the onset of neuronal differentiation. Cell Rep. 2022 Oct 25;41(4):111542. doi:10.1016/j.celrep.2022.111542 PubMed PMID: 36288718; PubMed Central PMCID: PMC9631114.

52. Michlewski G, Sanford JR, Cáceres JF. The Splicing Factor SF2/ASF Regulates Translation Initiation by Enhancing Phosphorylation of 4E-BP1. Molecular Cell. 2008 Apr 25;30(2):179–89. doi:10.1016/j.molcel.2008.03.013

53. Stutz F, Bachi A, Doerks T, Braun IC, Séraphin B, Wilm M, et al. REF, an evolutionary conserved family of hnRNP-like proteins, interacts with TAP/Mex67p and participates in mRNA nuclear export. RNA. 2000 Apr;6(4):638–50. doi:10.1017/s1355838200000078 PubMed PMID: 10786854; PubMed Central PMCID: PMC1369944.

54. Becker K, Bluhm A, Casas-Vila N, Dinges N, Dejung M, Sayols S, et al. Quantifying post-transcriptional regulation in the development of Drosophila melanogaster. Nat Commun. 2018 Nov 26;9(1):4970. doi:10.1038/s41467-018-07455-9

55. Ji Y, Tulin AV. Poly(ADP-Ribosyl)ation of hnRNP A1 Protein Controls Translational Repression in Drosophila. Molecular and Cellular Biology. 2016 Oct 1;36(19):2476–86. doi:10.1128/MCB.00207-16 PubMed PMID: 27402862.

56. Labourier E, Allemand E, Brand S, Fostier M, Tazi J, Bourbon HM. Recognition of exonic splicing enhancer sequences by the Drosophila splicing repressor RSF1. Nucleic Acids Res. 1999 Jun 1;27(11):2377–86. doi:10.1093/nar/27.11.2377

57. Goralski TJ, Edström JE, Baker BS. The sex determination locus transformer-2 of Drosophila encodes a polypeptide with similarity to RNA binding proteins. Cell. 1989 Mar 24;56(6):1011–8. doi:10.1016/0092-8674(89)90634-x PubMed PMID: 2493992.

58. Hilgers V. Regulation of neuronal RNA signatures by ELAV/Hu proteins. Wiley Interdisciplinary Reviews: RNA. 2023 Mar 1;14(2):e1733. doi:10.1002/wrna.1733

59. Temme C, Simonelig M, Wahle E. Deadenylation of mRNA by the CCR4–NOT complex in Drosophila: molecular and developmental aspects. Front Genet. 2014 May 26;5. doi:10.3389/fgene.2014.00143

60. Flanagan K, Baradaran-Heravi A, Yin Q, Dao Duc K, Spradling AC, Greenblatt EJ. FMRP-dependent production of large dosage-sensitive proteins is highly conserved. Genetics. 2022 Aug 1;221(4):iyac094. doi:10.1093/genetics/iyac094

61. Chen N, Zhang Y, Adel M, Kuklin EA, Reed ML, Mardovin JD, et al. Local translation provides the asymmetric distribution of CaMKII required for associative memory formation. Current Biology. 2022 Jun;32(12):2730–2738.e5. doi:10.1016/j.cub.2022.04.047

62. Kumar S, Lopez AJ. Negative feedback regulation among SR splicing factors encoded by Rbp1 and Rbp1-like in Drosophila. EMBO J. 2005 Jul 20;24(14):2646–55. doi:10.1038/sj.emboj.7600723 PubMed PMID: 15961996; PubMed Central PMCID: PMC1176452.

63. Parikh RY, Nayak D, Lin H, Gangaraju VK. Drosophila Modulo is essential for transposon silencing and developmental robustness. J Biol Chem. 2025 Mar;301(3):108210. doi:10.1016/j.jbc.2025.108210 PubMed PMID: 39848495; PubMed Central PMCID: PMC11879677.

64. Zheng W, Fong JHC, Wan YK, Chu AHY, Huang Y, Wong ASL, et al. Discovery of regulatory motifs in 5′ untranslated regions using interpretable multi-task learning models. Cell Systems. 2023 Dec 20;14(12):1103–1112.e6. doi:10.1016/j.cels.2023.10.011

65. He J, Xiong L, Shi S, Li C, Chen K, Fang Q, et al. Deep learning prediction of ribosome profiling with Translatomer reveals translational regulation and interprets disease variants. Nat Mach Intell. 2024 Nov;6(11):1314–29. doi:10.1038/s42256-024-00915-6

66. Huang L, Zhang H, Deng D, Zhao K, Liu K, Hendrix DA, et al. LinearFold: linear-time approximate RNA folding by 5’-to-3’ dynamic programming and beam search. Bioinformatics. 2019 Jul 15;35(14):i295–304. doi:10.1093/bioinformatics/btz375

67. Liaw R, Liang E, Nishihara R, Moritz P, Gonzalez JE, Stoica I. Tune: A Research Platform for Distributed Model Selection and Training [Internet]. arXiv; 2018 [cited 2025 Nov 7]. Available from: http://arxiv.org/abs/1807.05118 doi:10.48550/arXiv.1807.05118

68. Bergstra J, Yamins D, Cox D. Making a Science of Model Search: Hyperparameter Optimization in Hundreds of Dimensions for Vision Architectures. In: Proceedings of the 30th International Conference on Machine Learning [Internet]. PMLR; 2013 [cited 2025 Nov 7]. p. 115–23. Available from: https://proceedings.mlr.press/v28/bergstra13.html

69. Loshchilov I, Hutter F. Decoupled Weight Decay Regularization [Internet]. arXiv; 2019 [cited 2025 Nov 7]. Available from: http://arxiv.org/abs/1711.05101 doi:10.48550/arXiv.1711.05101

70. Attanasio G, Pastor E, Bonaventura CD, Nozza D. ferret: a Framework for Benchmarking Explainers on Transformers. In: Proceedings of the 17th Conference of the European Chapter of the Association for Computational Linguistics: System Demonstrations [Internet]. 2023 [cited 2025 Feb 13]. p. 256–66. Available from: http://arxiv.org/abs/2208.01575 doi:10.18653/v1/2023.eacl-demo.29

71. Virtanen P, Gommers R, Oliphant TE, Haberland M, Reddy T, Cournapeau D, et al. SciPy 1.0: fundamental algorithms for scientific computing in Python. Nat Methods. 2020 Mar;17(3):261–72. doi:10.1038/s41592-019-0686-2

72. Seabold S, Perktold J. Statsmodels: Econometric and Statistical Modeling with Python. SciPy 2010. 2010 May 1. doi:10.25080/Majora-92bf1922-011

73. Hunter JD. Matplotlib: A 2D Graphics Environment. Computing in Science & Engineering. 2007 May;9(3):90–5. doi:10.1109/MCSE.2007.55

74. Kurtzer GM, Sochat V, Bauer MW. Singularity: Scientific containers for mobility of compute. PLOS ONE. 2017 May 11;12(5):e0177459. doi:10.1371/journal.pone.0177459

75. Ray D, Kazan H, Cook KB, Weirauch MT, Najafabadi HS, Li X, et al. A compendium of RNA-binding motifs for decoding gene regulation. Nature. 2013 Jul 11;499(7457):172–7. doi:10.1038/nature12311 PubMed PMID: 23846655; PubMed Central PMCID: PMC3929597.

76. Mann HB, Whitney DR. On a Test of Whether one of Two Random Variables is Stochastically Larger than the Other. The Annals of Mathematical Statistics. 1947 Mar;18(1):50–60. doi:10.1214/aoms/1177730491

77. Benjamini Y, Hochberg Y. Controlling the False Discovery Rate: A Practical and Powerful Approach to Multiple Testing. Journal of the Royal Statistical Society: Series B (Methodological). 1995;57(1):289–300. doi:10.1111/j.2517-6161.1995.tb02031.x

78. RNA-binding profiles of Drosophila CPEB proteins Orb and Orb2 - PubMed [Internet]. [cited 2026 Apr 21]. Available from: https://pubmed.ncbi.nlm.nih.gov/27791065/

79. Yang YCT, Di C, Hu B, Zhou M, Liu Y, Song N, et al. CLIPdb: a CLIP-seq database for protein-RNA interactions. BMC Genomics. 2015 Feb 5;16(1):51. doi:10.1186/s12864-015-1273-2

